# Permissive aggregative group formation favors coexistence in yeast

**DOI:** 10.1101/2021.12.03.471114

**Authors:** Tom E. R. Belpaire, Jiří Pešek, Bram Lories, Kevin J. Verstrepen, Hans P. Steenackers, Herman Ramon, Bart Smeets

## Abstract

In *Saccharomyces cerevisiae*, the *FLO1* gene encodes flocculins that lead to formation of multicellular flocs, that offer protection to the constituent cells. Flo1p was found to preferentially bind to fellow cooperators compared to defectors lacking *FLO1* expression, resulting in enrichment of cooperators within the flocs. Given this dual function in cooperation and kin recognition, *FLO1* has been termed a ‘green beard gene’. Because of the heterophilic nature of Flo1p binding however, we hypothesize that kin recognition is permissive and depends on the relative stability of *FLO1*^+^/*flo1*^*−*^ versus *FLO1*^+^/*FLO1*^+^ bonds, which itself can be dependent on environmental conditions and intrinsic cell properties. We combine single cell measurements of adhesion strengths, individual cell-based simulations of cluster formation and evolution, and in vitro flocculation experiments to study the impact of relative bond stability on defector exclusion as well as benefit and stability of cooperation. We hereto vary the relative bond stability by changing the shear flow rate and the inherent bond strength. We identify a marked trade-off between both aspects of the green beard mechanism, with reduced relative bond stability leading to increased kin recognition, but at the expense of decreased cluster sizes and benefit of cooperation. Most notably, we show that the selection of *FLO1* cooperators is negative-frequency dependent, which we directly attribute to the permissive character of the Flo1p bond. Taking into account the costs associated to *FLO1* expression, this asymmetric selection results in a broad range of ecological conditions where coexistence between cooperators and defectors is stable. Although the kin recognition aspect of the *FLO1* ‘green beard gene’ is thus limited and condition dependent, the negative-frequency dependency of selection can conserve the diversity of flocculent and non-flocculent phenotypes ensuring flexibility towards variable selective pressures.

## Introduction

The transition towards multicellularity is one of the major developments that has driven the evolution of complex life^1–3^. Initially, independent individuals form facultative cooperative groups which can serve as a starting point for the evolution of obligate multicellular organisms wherein the individuals lose the ability to replicate independently^4–6^. These facultative groups can be formed through two distinct operations: formation of aggregative groups, known as ‘coming together’ (CT), and clonal growth, where the offspring remains closely associated with the parental cell, known as ‘staying together’ (ST)^4,7,8^. ST gives rise to clonal groups with high genetic relatedness whereas CT may also result in genetically mixed groups.

In featuring both unicellular lifestyles and various group phenotypes, *Saccharomyces cerevisiae* serves as a paradigm for studying ST^9–11^ and CT^12–14^ group formation, although *S. cerevisiae* does not have any known obligate multicellular de-scendants^4^. A key gene family involved in group formation in yeast comprises the *FLO* genes, which encode for flocculins, proteins involved in cell adhesion^4,12,15–19^. These flocculins possess an N-terminal domain protruding from the cell surface, a central domain of tandem repeated sequences, and a C-terminal glycosylphosphatidylinositol (GPI) domain anchored in the cell wall^15,20^. Based on the N-terminal domain, two types of flocculins can be distinguished. Flo11p harbors a fibronectin type III-like domain that confers homophilic protein-protein interaction with neighbouring cells^11,21^. Flo11p-mediated ad-hesion partakes in multiple ST group phenotypes such as biofilm^11,22^ and pseudohyphae formation^23^. In contrast, the *FLO1* gene encodes for a PA14-like N-terminal domain that binds to mannose residues on the cell wall of neighbouring cells, a mechanism that is heterophilic in nature^24,25^. Flo1p controls the CT flocculation phenotype, causing yeast cells to aggregate and form flocs in agitated suspensions. When sufficiently large, these flocs offer protection to the constituent cells against chemical^12^ and biological^26,27^ stress. Furthermore, flocs ensure rapid sedimentation to escape undesirable conditions^28^. As such, floc formation is a type of cooperative behavior in which the benefit only exists when sufficient individuals participate^12^.

In addition to facilitating group formation, both types of *FLO* genes also permit kin recognition through selective adhesion. In this quality, they have been identified as ‘green beard genes’, a single set of alleles that promotes cooperation while also excluding non-collaborating individuals (defectors)^12,29,30^. In case of *FLO11* selective adhesion is mediated by the homophilic nature of the interaction^11^. Flo1p was also found to preferentially bind to fellow cooperators compared to defectors, resulting in enrichment of cooperators within the flocs^12^. The heterophilic nature of the Flo1p bond however also permits adhesion to non-producer cells. The observed enrichment of cooperator cells might then be explained by a higher bond strength between cooperators due to the potential of reciprocity in homotypic *FLO1*^+^/*FLO1*^+^ interactions. We hypothesize that kin recognition via such heterophilic binding is however only partially selective and dependent on the relative stability of *FLO1*^+^/*flo1*^*−*^ versus *FLO1*^+^/*FLO1*^+^ bonds, which itself depends on the intrinsic bond properties but also the tensile forces trying to separate interacting cells. Since *S. cerevisiae* lacks intrinsic motility, shear forces arising due to fluid flow are thought to be the main instigators of bond formation and breakage events. Because of this potentially ‘permissive’ nature of the *FLO1* kin recognition, defectors might still be able to invade flocs and exploit the benefits of cooperation^11,14^. Since defectors do not pay the metabolic cost associated with Flo1p production, they can have an increased fitness relative to the cooperators. Consequently, the evolutionary stability of *FLO1* is not guaranteed. Knowing the impact of relative bond stability and its driving factors on kin recognition is therefore critical to understand the evolution of CT flocculation driven by heterophilic Flo1p adhesion.

In this work, we evaluate the impact of the relative bond stability of heterotypic and homotypic Flo1p interactions on the exclusion of defectors and the evolutionary stability of flocculation in mixed populations with varying cooperator frequencies. To this end, we first determine the intrinsic relative stability of heterotypic and homotypic interactions using single cell-force spectroscopy (SCFS) and subsequently characterize the extent of permissiveness in Flo1p-mediated kin recognition. Based on these measured bond properties, we evaluate both the cooperative benefits and the degree of kin recognition of the *FLO1* green beard cooperation and its evolutionary stability using cell-based simulation of shear-induced CT group formation. We conclude that the relative stability of heterotypic and homotypic interactions, modulated by varying either tensile shear stresses or bond properties, determines a trade-off between kin recognition and cooperative benefits. Remarkably, size-dependent selection of clusters results in a negative-frequency-dependent selection pressure that stabilizes coexistence between defectors and cooperators in a broad range of ecological and mechanical conditions. Stable coexistence ensures the retention of diversity and thus facultative group formation, which might eventually give rise to evolution of obligate multicellularity.

## Materials and Methods

### Yeast strains and media

All yeast strains used are listed in Table 1. Yeast cultures were first cultured in YPD for 3 days and subsequently inoculated in YPG and grown for 2 days. Aftwards, cells were harvested and washed once in 200 mM EDTA and twice in milliQ.

### Single-cell force spectroscopy

Single-cell force spectroscopy was performed as described by^24^. In short, cell probes were prepared by immobilizing single yeast cell on polydopamine-functionalized tipless cantilevers. The cell probe was brought into contact with single cells immobilized on a glass coverslip with polydopamine using a maximum contact force of 1 nN, retract velocity of 1 *μ*m/s and contact time of 1 s in the presence of 200 *μ*M CaCl_2_. Cell viability of both the cell probe and the immobilized cells on the substrate were followed by the FUN-1 cell stain throughout the measurement.

### Flocculation assays

After harvesting and washing yeast cells at various ratios of *FLO1*^+^ and *flo1*^*−*^, cells were inoculated in 5ml milliQ with a final density of 3.0±1.4 10^6^ cells/mL. After inoculation the tubes were carefully turned to homogenize them and sampled for the initial ratio of cells *x*_*i*_. Test tubes were shaken on an orbital shaker at varying agitation rates (0, 100, 200 or 400 RPM) for 5 min. After agitation, the flocs were allowed to settle for 5 min after which the sedimented fraction was sampled *x*_out_. Prior to cell counting using flow cytometry, samples were washed with 200 mM EDTA to disrupt any floc formation. Ratios *x* were determined as the fraction of red *FLO1*^+^ cells versus the total amount of cells. The experiments were performed in 10 mM CaCl_2_ necessary for flocculation, and in milliQ as a control.

### Individual cell-based model

We performed simulations of a center-based cell model in an overdamped system in laminar flow with periodic boundary conditions. External shear force is imposed based on a set shear rate 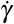 Cell-cell interaction was modeled using a linear adhesion model with rupture force *F*_*d*_ and rupture distance *d*_*r*_, Hertzian repulsion and linear intercellular viscosity. Cell velocities were computed by solving 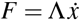, with Λ the combined friction/resistance matrix. Positions are updated according to the explicit Euler method. A full description of the computational methods is given in the SI text.

## Results

### Flo1p confers heterophilic cell-cell adhesion

Flo1p flocculins bind to mannose residues on neighbouring cells. To quantify the force resulting from these adhesive bonds, we employ the SCFS method described by El-Kirat-Chatel et al.^24^. We test three types of interaction pairs: *flo1*^*−*^/*flo1*^*−*^, *FLO1*^+^/*flo1*^*−*^ and *FLO1*^+^/*FLO1*^+^. For every interaction type, we measure the detachment force *F*_*d*_ and the rupture distance *d*_*r*_ of the bond (Fig. 1 A-F). We find that the detachment force *F*_+*−*_ of the heterotypic interaction is approximately half (*≈* 55%) of the homotypic *FLO1*^+^ interaction, whereas a homotypic *flo1*^*−*^ interaction is an order of magnitude (*≈* 7%) lower in adhesive strength. This bond heterophilicity is consistent with a permissive kin recognition mechanism^14^. The differential adhesion hypothesis (DAH) predicts three different modes of group organization based on the ratio of interaction energy between heterotypic and homotypic bonds; segregation, spreading of the weakly adhering cell type, and intermixing of both cell types^31^. For the bond energy associated with the measured detachment force and rupture distance of Flo1p, the DAH predicts spreading for the majority of observations (Fig. 1G-H, Fig S1). Spreading of *flo1*^*−*^ cells around a central *FLO1*^+^ cluster has previously been observed and produces additional benefits in macroscopic flocs. For example, in the case of protection against antifungal compounds such as amphotericin B, the outer layer of *flo1*^*−*^ cells can serve as ‘living shield’, leading to increased protection of the *FLO1*^+^ cells at the core of the floc^12^. In contrast, the absence of segregation indicates potential for exploitation of the flocculation by the *flo1*^*−*^ cells, as segregation is thought to be the ideal scenario for cooperative phenotypes^32–34^. The DAH predicts the equilibrium configuration of a mixture of cells with differing interaction energy. However, yeast cells are too large to be significantly agitated by thermal forces, and they lack intrinsic cell motility. Consequently, in real-life conditions, substantial energy barriers can prevent the system from relaxing to its equilibrium configuration. Hence, to evaluate the degree of kin recognition due to *FLO1* expression, the driving forces responsible for floc formation must be taken into account.

**Figure 1.**
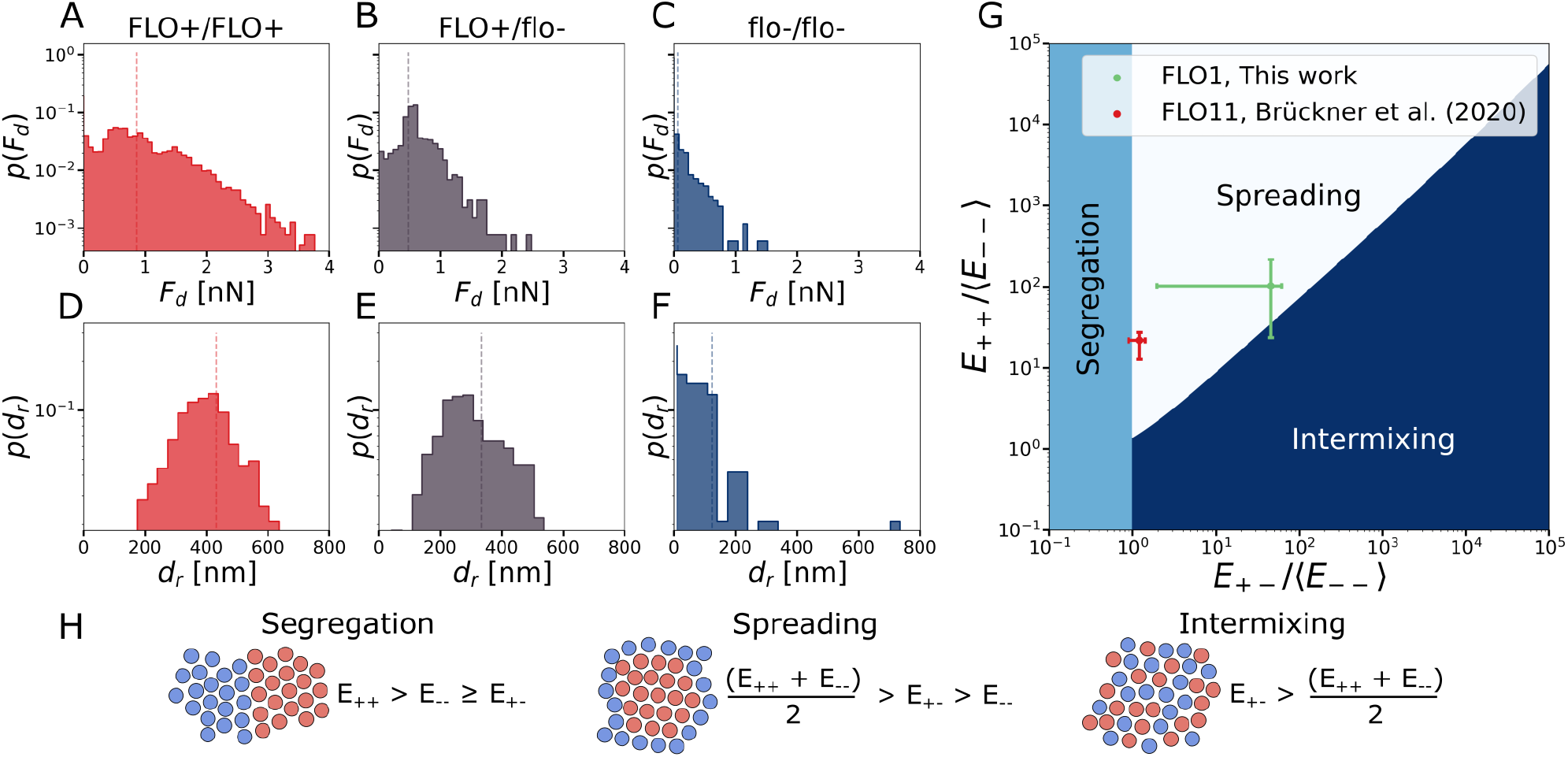
Mechanical measurement of Flo1p bond properties and mixing predictions. (A-C) Probability density functions of the measured maximum detachment force *F*_*d*_ for a *FLO1*^+^/*FLO1*^+^ (A), *FLO1*^+^/*flo1*^*−*^ (B) and *flo1*^*−*^/*flo1*^*−*^ (C) interaction. (D-F) Probability density function of the rupture length *d*_*r*_ of the bonds, measured by maximum distance with significant adhesive forces. (G) Based on the bond energies, *E* = *F*_*d*_*d*_*r*_*/*2, the colony structure was predicted by the differential adhesion theory (DAH). Single cell-force spectroscopy data of Flo11p was obtained from^11^. (H) DAH predicts segregation, spreading and intermixing based on the ratio of bond energies.

### Shear flow promotes relatedness in mixed clusters at the expense of cooperative benefits

Flocs originate from collisions between individual yeast cells, which are facilitated by external forces, such as shear flow. In practice, the formation of large flocs is realized in two stages: 1) nucleation and growth of small clusters due to collisions in shear flow and 2) differential sedimentation and size-based separation of clusters, leading to macroscopic flocs. We evaluate the size and composition of cell clusters in a minimal linear shear simulation with varying initial cooperator frequency *x*_*i*_ and shear rate 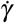 (Fig. 2A-E). Since the composition of large flocs is chiefly determined by clusters that are of sufficient size to sediment, it is reasonable to assume that the main benefit of cooperation is increased cluster size, an effect also observed in other model systems^7,9,26,35,36^. At sufficient cell density, the average cluster size *C* shows an exponential shear-dependent relaxation over time towards a dynamic steady-state. In contrast, at low density, a slowed down relaxation is observed, indicative of granular compaction in between infrequent collision events (Fig. 2F, S2). In both density regimens, the steady-state cluster size *C*_∞_ decreases with shear rate. Moreover, cluster size increases with the initial fraction of cooperators (Fig. 2G). Overall, enrichment of *FLO1*^+^ cells is observed in clusters, which increases with shear rate as heterotypic bonds become unstable at lower shear rates than homotypic bonds (Fig. 2H, S6). However, due to the permissive binding mechanism, selection for *FLO1*^+^ is weak. This is further apparent in the relatedness 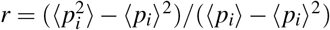, with *p*_*i*_ the fraction of cooperators in cluster *i*, which is thus evaluated at the level of whole clusters rather than the level of single cells and their direct neighbours. Relatedness signifies the directness of cooperative interactions, with *r* = 1 in populations with clusters uniquely composed of *FLO1*^+^ or *flo1*^*−*^ cells and *r* = 0 in absence of variation in cluster composition^37^. In general, we find only modest relatedness (*r* < 0.3) in all shear regimens, characteristic for CT group formation and permissive kin recognition^37,38^. Nonetheless, relatedness is favored by increasing shear rate (Fig. 2I, S6). Finally, we also observe radial assortment within clusters, as was noted by Smukalla *et al*.^12^ in much larger flocs, and in line with the equilibrium conditions predicted by the DAH (Fig. S7). However, this assortment is not very pronounced, and these micro-assorted clusters are too small to provide protection to realistic chemical stress conditions. In conclusion, shear-driven aggregation of mixtures of cooperators and defectors leads to partially selective group formation due to the exclusion of defectors from clusters, which results in smaller but more selective clusters with increasing shear rate. However, selection is not very efficient due to the permissive binding mechanism of Flo1p that allows for a heterogeneous cluster composition at all shear conditions, and is markedly different from the thermodynamical equilibrium predicted by DAH (Fig. 1G).

**Figure 2.**
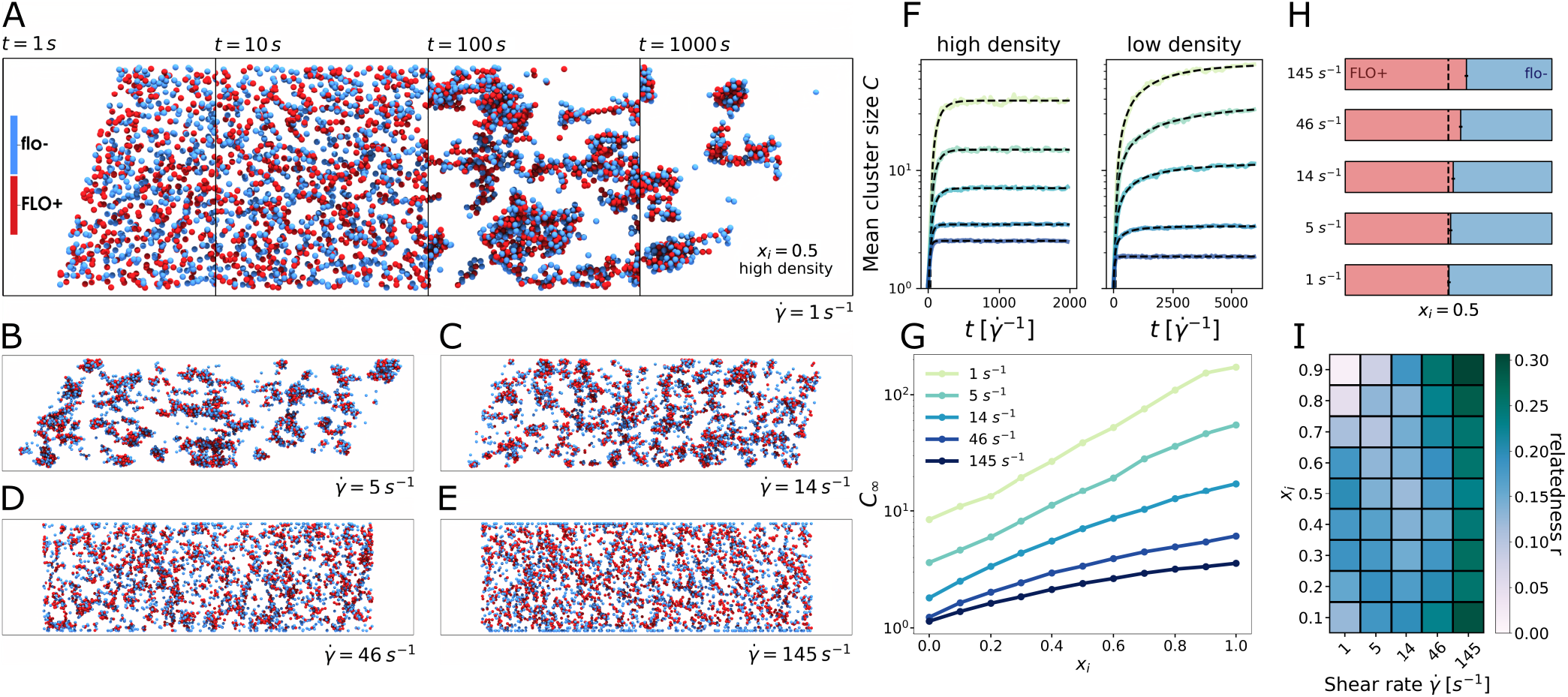
Effect of shear on heterotypic Flo1p-dependent flocculation. (A) Temporal progression of flocculation starting from a homogeneously mixed population of *FLO1*^+^ (red) and *flo1*^*−*^ (blue) at increasing time points 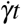, shown for a cooperator frequency *x*_*i*_ = 0.5, high density, *ρ*_high_ = 1.66 10^7^ cells/ml and shear rate 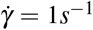.(B-E) Endpoint of flocculation at various shear rates, shown for cooperator frequency *x*_*i*_ = 0.5. (F) Time evolution of the mean cluster size *C* for high (*ρ*_high_ = 1.66 *×* 10^7^ cells/ml) and low density (*ρ*_low_ = 0.83 *×* 10^7^ cells/ml) for varying shear rate. The black lines indicate exponential fit *C*(*t*) = *C*_∞_ [1*−*exp(*−t/τ*)] and a stretched exponential *C*(*t*) = *C*_∞_[1*−*exp(*−* (*t/τ*)^*β*^)] fit for the high and the low density respectively. At high (‘super-critical’) density, the projected area, integrated across a circular flow line is larger than one, and the system reaches a dynamic steady-state. At low (‘sub-critical’) density, this projected area is lower than one, and collisions become exceedingly rare after closed flow lines have been depleted of cells, see Fig. S2,S3. (G) Steady state cluster size *C*_∞_ in function of cooperator frequency *x*_*i*_ for varying shear rate, see Fig. S4. (H) Cluster composition for clusters of size *>* 2 cells for varying shear rate. The dotted black line indicates cooperator frequency *x*_*i*_ = 0.5. (I) Cluster relatedness in function of shear rate 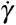 and cooperator frequency *x*_*i*_, see Fig. S6.

### Permissive kin recognition facilitates coexistence

The evolutionary robustness of *FLO1* depends on its associated costs and benefits. We evaluate both by using a conceptual modeling framework consisting of three sequential ecological processes (Fig. 3A). First, a mixed population with cooperator frequency *x*_*i*_ is exposed to shear flow and allowed to flocculate (Fig. 2). Second, individual cells are selected based on the steady-state size and thus the expected benefit offered by the cluster they belong to, *C*_∞_. The probability to survive is 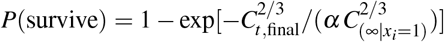, with 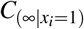 the mean cluster size for a fully cooperative system and *α* a parameter tuning the selection strength. *P*(survive) is based on the Stokesian sedimentation velocity 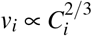, i.e. larger clusters sediment faster and have an increased selection probability (Fig. 3B, S9). Third, the selected cells are allowed to exponentially grow for a number of generations^39,40^, taking into account a 3% percent fitness deficit for the *FLO1*^+^ cells relative to the *flo1*^*−*^ cells (Fig. 3C)^12^. After flocculation, selection and growth, the population drift Δ*x*_*i*_ is determined based on the frequency of cooperators before and after each of the three steps, Δ*x*_*i*_ = *x*_out_ *−x*_*i*_.

**Figure 3.**
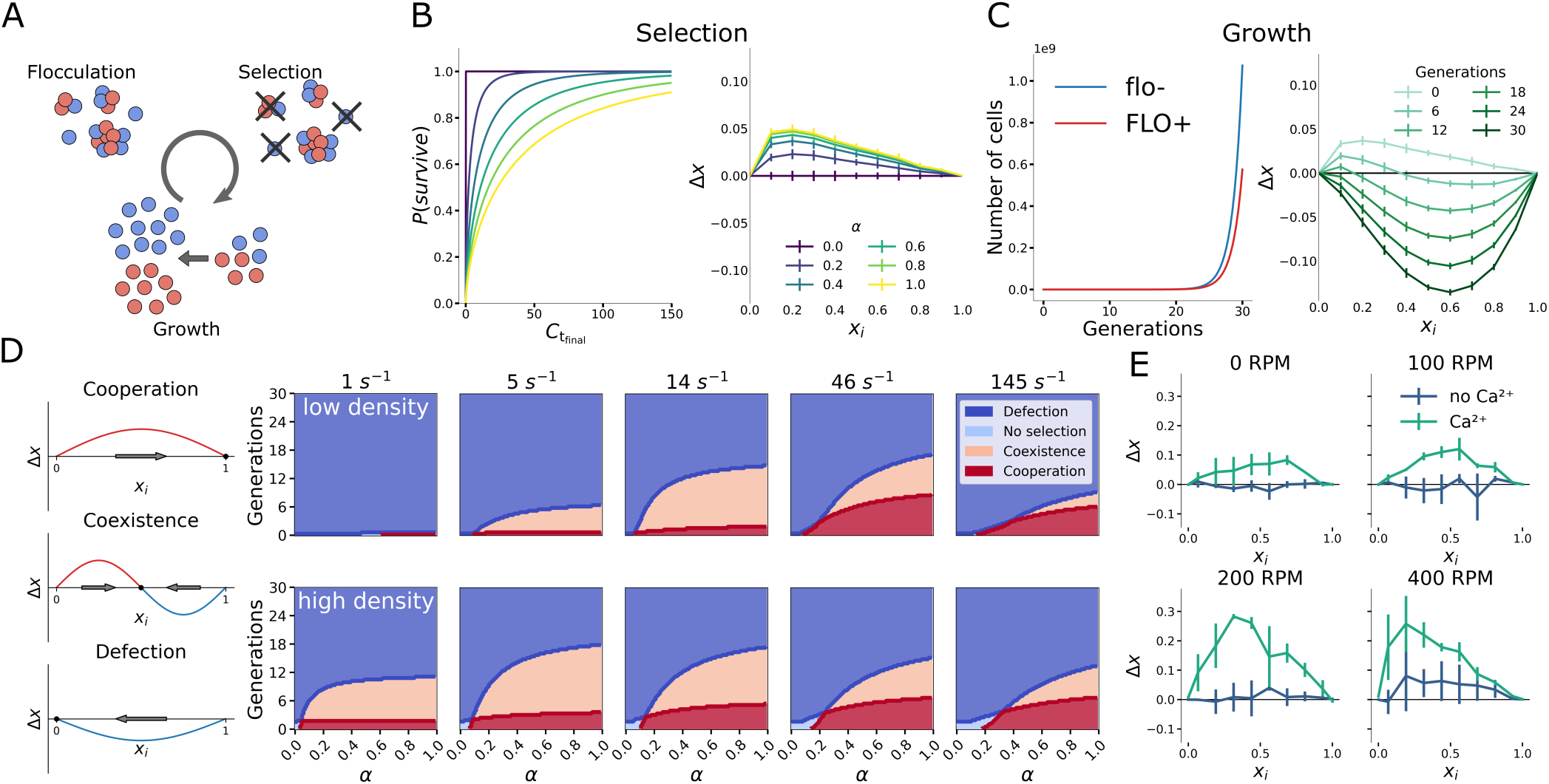
Population dynamics in *FLO1* cooperation. (A) Three sequential ecological processes are considered; flocculation, selection by sedimentation and growth. (B) Cluster size selection probability *P*(survive) in function of steady-state cluster size *C*_*t*,final_. Population drift Δ*x* after selection at varying selection strength *α* is shown for 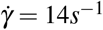. (C) selected *FLO1*^+^ cells experience a fitness deficit relative to *flo1*^*−*^ of 3% as reported by Smukalla *et al*.^12^. Drift curves at moderate selection strength 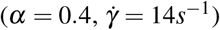 and increasing generations. (D) Classification of drift curves in evolutionarily stable strategies (ESS) cooperation, coexistence, defection. ESS in function of *α* and the number of generations for high and low density, see Fig. 2. (E) Experimental characterization of population drift for various rotor amplitudes in the presence and absence of Ca^2+^.

After flocculation and selection (thus prior to the growth step), there is a preferential retention of cooperating cells for *α >* 0. Markedly, the peak in population drift after selection showcases an asymmetry towards a lower cooperator frequency (Fig. 3B, S10). At low *x*_*i*_, only clusters with a frequency of cooperators ≫*x*_*i*_ are sufficiently strong to resist the disruptive force from shear flow. This results in a relative enrichment of cooperators in the surviving clusters. Conversely, at large *x*_*i*_, the abundance of cooperators in clusters provides sufficient favorable locations for defector cells to be incorporated and the frequency of cooperators in clusters approaches the initial population frequency. Upon imposing a growth-associated cost for cooperation, this asymmetry can result in selection in favor of cooperators at low *x*_*i*_ (Δ*x >* 0) and selection for defectors at high *x*_*i*_ (Δ*x <* 0) (Fig. 3C). Based on the shape of the population drift curve, we determine the evolutionarily stable strategy (ESS) as a function of selection strength *α* (∝ social benefits) and number of growth generations (∝ social cost), and this at varying shear rate (Fig. 3D). Cooperation emerges as an ESS for increasing strength *α*. However, given a high number of generations — or high growth-associated costs — the resulting ESS is defection. This highlights that permissive kin recognition with permissive bonds is not efficient at fully excluding defector cells from cooperative groups without additional external selection pressure^14^. However, due to the aforementioned asymmetry in selection, coexistence is the ESS for a large range of ecological parameters. The stable point (i.e., the stable frequency of cooperators) shifts towards a lower cooperator frequency with higher number of generations (Fig. S11). Whereas cooperation is more favored with increasing shear rate, coexistence is notably favored at intermediate shear rate, where the asymmetry in selection is most pronounced (Fig.S10). Here, cooperative homotypic bonds are always stable, whereas permissive heterotypic bonds can be broken by tensile shear forces, thereby maximizing the relative enrichment of cooperators at low *x*_*i*_. Finally, at low cell density, clusters are more compact and collide less frequently compared to high cell density. Consequently, the peak in coexistence shifts towards higher shear rate, as more shear force is required to penalize the incorporation of permissive bonds in dense, well-connected clusters (Fig. S2).

For *in vitro* verification of the predicted asymmetric population drift, we mimic the first two steps of the evolutionary framework, flocculation and selection, using a simple flocculation-sedimentation assay, where we inoculate various cooperator fractions and agitate them at varying rotator speed. Selection is performed by sampling from the sediment, which contains flocs that preferentially consist of larger clusters (Fig. S12). Based on the frequency of cooperators in the sedimented (i.e., selected) flocs and the inoculum frequency, the drift was estimated (Fig. 3E). Increasing the rotor speed (∝ shear rate) resulted in increasingly positive drift curves in the presence of Ca^2+^ — which is required for flocculation — indicating an increased exclusion of *flo1*^*−*^ cells, as observed *in silico* (Fig. 2). In addition, at sufficient rotor speed (200 RPM and 400 RPM) the maximal drift is located at a lower cooperator frequency, demonstrating the same characteristic asymmetry in *FLO1*^+^ enrichment that was predicted from *in silico* simulations (Fig. 3B). When including a fixed cost incurred by growth, these drift curves will give rise to coexistence as an ESS for a mixed population of *FLO1*^+^ and *flo1*^*−*^ cells (Fig. 3D).

### Flo1p bond mechanism permits evolutionary flexibility

The emergence of coexistence due to asymmetric selection is contingent on permissive interactions and is absent in a hypothetical scenario with direct kin recognition where *F*_+*−*_ = *F*_*−−*_. In case of direct kin recognition, symmetrical drift expands the fully cooperative region at the expense of coexistence (Fig. 4A-B). Due to complete exclusion of defector cells from clusters, cooperation is stable irrespective of the number of generations and the *FLO1*-associated costs, given sufficient selective strength (*α >* 0.4) (Fig. 4B-C). Remarkably, complete exclusion of defector cells in direct kin recognition also results in smaller clusters, and thus cooperation-associated benefits, in mixed populations (0 *< x*_*i*_ *<* 1) (Fig. 4D). Moreover, since the cooperation-associated benefits are small at low *x*_*i*_, the emergence of direct kin recognition, e.g. by mutation of a single cell, is not expected to perpetuate in an initially fully defective population (*x*_*i*_ = 0) in CT group formation. In contrast, permissive recognition is more favorable to develop due to higher cooperation-associated benefits and asymmetric population drift. In exchange for the resistance to defection provided by the increased selectivity of direct kin recognition, the permissiveness of the Flo1p bond permits evolutionarily stable flexibility, by conserving coexistence between cooperators and defectors.

**Figure 4.**
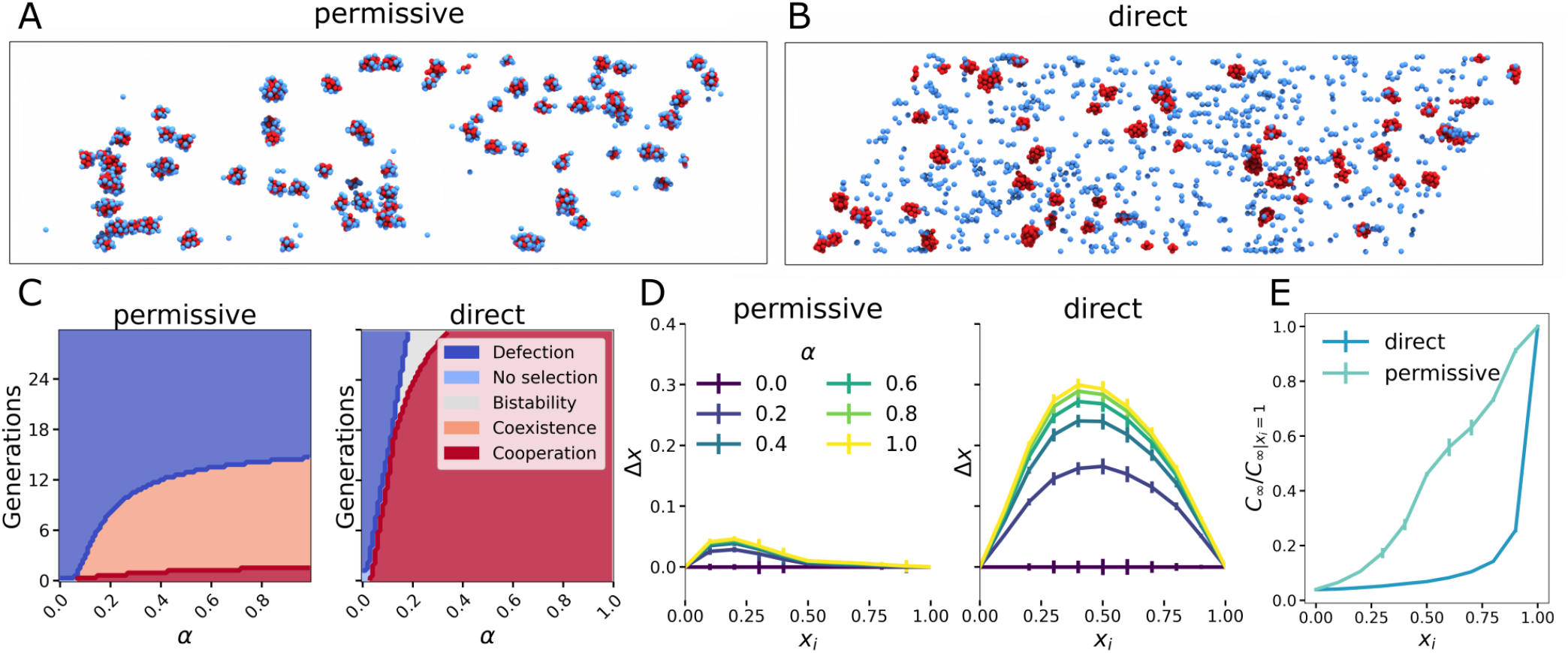
Effect of Flo1p bond properties on evolutionary stability. (A) Evolutionary drift Δ*x* after flocculation and selection for permissive and direct kin recognition. (B) Evolutionarily stable strategy (ESS) for permissive and direct kin recognition. For direct kin recognition, bistability emerges when Δ*x* is increasing at the zero point^47^. (C) Relatedness *r* in function of initial cooperator frequency *x*_*i*_ for permissive and direct kin selection. (D) Cooperative benefits relative to the fully cooperative system *x*_*i*_ = 1 for both permissive and direct kin recognition. (E) Empirical *FLO1*^+^/*FLO1*^+^ detachment force variability *F*_*d*_. Effect of bond strength on the ESS at strong selection (*α* = 1). (F) Final cluster size 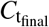 in function of initial cooperator frequency *x*_*i*_ for varying homotypic detachment forces *F*_++_, conserving *F*_++_ *≈* 2*F*_+*−*_. (G) Relatedness *r* in function of *F*_++_, conserving *F*_++_ *≈* 2*F*_+*−*_ shown for *x*_*i*_ = 0.5. Results are shown for low density (*ρ*_low_ = 0.83 *×* 10^7^ cells/ml) and shear rate 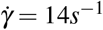.

In contrast to the hypothetical nature of direct kin recognition due to the Flo1p bond mechanism, variability in adhesive strength in flocculation has been observed to arise due to stochasticity in bond formation (Fig. 1A-C, Fig. 4E), or variation in the intragenic tandem repeats of *FLO1*, which are known to undergo frequent recombination events^18,24^. Varying the homotypic adhesive strength *F*_++_ while conserving the ratio *F*_++_ *≈* 2*F*_+−_ highlights the dilemma of a permissive green beard gene: In case of an increase in adhesion, the cooperative benefits increase (Fig. 4F), but this weakens kin recognition due to the increased stability of the heterotypic bond (Fig. 4G). As such, increased homotypic adhesive strength expands the stability of coexistence at the expense of cooperation (Fig. 4E). Furthermore, the adhesive force at which cooperation is maximally stable depends on the shear rate. This provides a possible explanation for the great variability in tandem repeats of *FLO1*, as it allows flexibility in the aggregative strategy to adapt to heterogeneous environments.

## Discussion

*FLO1* has been identified as a green beard gene governing both aggregation and kin recognition during flocculation^12^. Here, we provide evidence that reciprocity in purely cooperative (homotypic *FLO1*^+^/*FLO1*^+^) interactions is associated with increased detachment force compared to exploitative (heterotypic *FLO1*^+^/*flo1*^*−*^) interactions. However, as cooperators are still vulnerable to exploitative interactions, the kin recognition mechanism of Flo1p is permissive and only weakly directs the cooperative benefits to Flo1p-producing individuals. This is in marked contrast with *FLO11*, which confers homophilic adhesion that leads to direct kin recognition and has been implied in sub-species level discrimination based on a single genetic difference^11^. Our results indicate that varying the relative bond stability of cooperative and exploitative interactions can modulate between both facets of the *FLO1* green beard mechanism: kin recognition and cooperative benefits. We explore shear flow and bond strength, respectively an environmental and intrinsic factor affecting relative bond stability. First, at low shear rate, both cooperative and exploitative interactions are stable resulting in large clusters (∝ cooperative benefits) with low relatedness (∝ kin recognition). High shear rate primarily leads to instability of the exploitative interaction, resulting in smaller clusters but with increased relatedness. Second, the high mobility of tandem repeated sequences of the *FLO1* gene is thought to modulate the adhesive forces between cells^24^ and has been shown to result in phenotypic heterogeneity^18^. Assuming the generality of *F*_++_*≈*2*F*_+*−*_, we predict that increasing *FLO1* gene length, and consequently adhesive forces, results in greater cooperative benefits but weaker kin recognition at a given shear rate. We propose that high variability of *FLO1* gene length allows adaptation towards the more appropriate strategy, increasing kin recognition in weak selective regimens or increasing benefits in stringent selection, and thereby potentially stabilizes flocculation in changing environments.

For permissive kin recognition due to the heterophilic nature of Flo1p, we predict a negative-frequency-dependent selection (NFDS) in function of the cooperator frequency. NFDS arises when a decrease in cooperator frequency more severely disadvantages defectors^41^. In our case, decreasing cooperator frequency decreases the probability of defector incorporation in the clusters at favorable locations and decreases the stability of clusters with relatively high defector fractions. This results in a relative increase of cooperator enrichment at low cooperator frequency. In addition, we show that in case of homophillic interactions, and thus in the absence of permissiveness in kin recognition, NFDS is lost. As NFDS is a known driver of biological diversity^42,43^, it can stabilize the evolution of cooperative phenotypes^41,44^. In case of permissive kin recognition, we find a stable coexistence of cooperators and defectors in a wide range of cooperation-associated costs and benefits. Coexistence offers flexibility through diversity in environments that are characterized by transient and variable selection pressures, where permissive coming together group formation is thought to outperform staying together^14^. Furthermore, stable coexistence also permits the conservation of variability in *FLO1* gene length, stabilizing the aforementioned adaptability to the environment. Moreover, we postulate that due to the negative-frequency-dependency and the higher return of cooperative benefits, permissive kin recognition is more likely to emerge than direct kin recognition where contacts predominantly originate from stochastic collisions such as low nutrient environments. However, this also renders permissive kin recognition more prone to invasion of defectors.

Our results indicate that permissive CT group formation is susceptible to invasion of non-flocculent phenotypes and can conserve the diversity of a population. Coexistence implies within-group social conflict and is therefore believed to limit the direct further evolution of obligate multicellularity and its accompanied potential of complexity^45^. Nevertheless, we propose that this conserved diversity can facilitate the further evolution of different group formation phenotypes. As such, ST group formation has been shown to emerge in flocculating yeast populations and to synergistically improve population fitness^13,14^. On longer evolutionary timescales, emerging ST group formation has been shown to be able to outperform CT by flocculation overcoming aforementioned within-group social conflict^14,46^. Finally, we propose that the physical environment can modulate the significance of permissive CT group formation, thereby shaping the intricate balance between CT and ST, which are fundamental biological operations that can prompt complex biological construction respectively through specialization in obligate multicellularity or conservation of diversity^8,14^.

## Acknowledgements

The authors thank Karin Voordeckers for providing us with the yeast strains; Carmen Bartic and Olivier Deschaume for expertise and support with the SCFS experiments. This work was supported by the KU Leuven Research Fund (CELSA/18/031,C24/18/046). B.S. acknowledges support from the Research Foundation Flanders (FWO) grant 12Z6118N. Research in the lab of K.J.V. is supported by KU Leuven, Vlaams Instituut voor Biotechnologie (VIB) and FWO.

## Author contributions statement

H.R. and B.S. conceived the project, T.B., J.P., and B.S. designed and conducted the simulations. T.B. designed and conducted the experiments. T.B. and B.S. performed data analysis. T.B., H.S. and B.S. wrote the manuscript. All authors reviewed the manuscript.

## References

1. Szathmáry, E. Toward major evolutionary transitions theory 2.0. Proc. Natl. Acad. Sci. United States Am. 112, 10104–10111, 10.1073/pnas.1421398112 (2015).

2. Niklas, K. J. & Newman, S. A. The origins of multicellular organisms. Evol. Dev. 15, 41–52, 10.1111/ede.12013 (2013).

3. Pfeiffer, T. & Bonhoeffer, S. An evolutionary scenario for the transition to undifferentiated multicellularity. Proc. Natl. Acad. Sci. United States Am. 100, 1095–1098, 10.1073/pnas.0335420100 (2003).

4. Fisher, R. M. & Regenberg, B. Multicellular group formation in Saccharomyces cerevisiae. Proc. Royal Soc. B: Biol. Sci. 286, 10.1098/rspb.2019.1098 (2019).

5. Umen, J. G. Green Algae and the Origins of Multicellularity in the Plant Kingdom. Cold Spring Harb. Perspectives Biol. 6, a016170, 10.1101/cshperspect.a016170 (2014).

6. Knoll, A. H. The multiple origins of complex multicellularity. Annu. Rev. Earth Planet. Sci. 39, 217–239, 10.1146/annurev.earth.031208.100209 (2011).

7. Bonner, J. T. The origins of multicellularity. Integr. Biol. Issues, News, Rev. 1, 27–36, 10.1002/(sici)1520-6602(1998)1:1<27::aid-inbi4>3.3.co;2-y (1998).

8. Tarnita, C. E., Taubes, C. H. & Nowak, M. A. Evolutionary construction by staying together and coming together. J. Theor. Biol. 320, 10–22, 10.1016/j.jtbi.2012.11.022 (2013).

9. Ratcliff, W. C., Denison, R. F., Borrello, M. & Travisano, M. Experimental evolution of multicellularity. Proc. Natl. Acad. Sci. United States Am. 109, 1595–1600, 10.1073/pnas.1115323109 (2012).

10. Koschwanez, J. H., Foster, K. R. & Murray, A. W. Sucrose utilization in budding yeast as a model for the origin of undifferentiated multicellularity. PLoS Biol. 9, e1001122, 10.1371/journal.pbio.1001122 (2011).

11. Brückner, S. et al. Kin discrimination in social yeast is mediated by cell surface receptors of the flo11 adhesin family. eLife 9, 10.7554/eLife.55587 (2020).

12. Smukalla, S. et al. FLO1 Is a Variable Green Beard Gene that Drives Biofilm-like Cooperation in Budding Yeast. Cell 135, 726–737, 10.1016/j.cell.2008.09.037 (2008).

13. Driscoll, W. W. & Travisano, M. Synergistic cooperation promotes multicellular performance and unicellular free-rider persistence. Nat. Commun. 8, 10.1038/ncomms15707 (2017).

14. Pentz, J. T. et al. Ecological Advantages and Evolutionary Limitations of Aggregative Multicellular Development. Curr. Biol. 30, 4155–4164.e6, 10.1016/j.cub.2020.08.006 (2020).

15. Goossens, K. & Willaert, R. Flocculation protein structure and cell-cell adhesion mechanism in Saccharomyces cerevisiae. Biotechnol. Lett. 32, 1571–1585, 10.1007/s10529-010-0352-3 (2010).

16. Di Gianvito, P., Tesnière, C., Suzzi, G., Blondin, B. & Tofalo, R. FLO5 gene controls flocculation phenotype and adhesive properties in a Saccharomyces cerevisiae sparkling wine strain. Sci. Reports 7, 1–12, 10.1038/s41598-017-09990-9 (2017).

17. Veelders, M. et al. Structural basis of flocculin-mediated social behavior in yeast. Proc. Natl. Acad. Sci. United States Am. 107, 22511–22516, 10.1073/pnas.1013210108 (2010).

18. Verstrepen, K. J., Jansen, A., Lewitter, F. & Fink, G. R. Intragenic tandem repeats generate functional variability. Nat. Genet. 37, 986–990, 10.1038/ng1618 (2005).

19. Verstrepen, K. J. & Klis, F. M. Flocculation, adhesion and biofilm formation in yeasts. Mol. Microbiol. 60, 5–15, 10.1111/j.1365-2958.2006.05072.x (2006).

20. Verstrepen, K. J., Reynolds, T. B. & Fink, G. R. Origins of variation in the fungal cell surface. Nat. Rev. Microbiol. 2, 533–540, 10.1038/nrmicro927 (2004).

21. Kraushaar, T. et al. Interactions by the fungal Flo11 adhesin depend on a fibronectin type III-like adhesin domain girdled by aromatic bands. Structure 23, 1005–1017, 10.1016/j.str.2015.03.021 (2015).

22. Oppler, Z. J., Parrish, M. E. & Murphy, H. A. Variation at an adhesin locus suggests sociality in natural populations of the yeast saccharomyces cerevisiae. Proc. Royal Soc. B: Biol. Sci. 286, 10.1098/rspb.2019.1948 (2019).

23. Lo, W. S. & Dranginis, A. M. The cell surface flocculin Flo11 is required for pseudohyphae formation and invasion by Saccharomyces cerevisiae. Mol. Biol. Cell 9, 161–171, 10.1091/mbc.9.1.161 (1998).

24. El-Kirat-Chatel, S. et al. Forces in yeast flocculation. Nanoscale 7, 1760–1767, 10.1039/c4nr06315e (2015).

25. Kobayashi, O., Hayashi, N., Kuroki, R. & Sone, H. Region of Flo1 proteins responsible for sugar recognition. J. Bacteriol. 180, 6503–6510, 10.1128/jb.180.24.6503-6510.1998 (1998).

26. Kapsetaki, S. E. & West, S. A. The costs and benefits of multicellular group formation in algae*. Evolution 73, 1296–1308, 10.1111/evo.13712 (2019).

27. Quintero-Galvis, J. F. et al. Exploring the evolution of multicellularity in Saccharomyces cerevisiae under bacteria environment: An experimental phylogenetics approach. Ecol. Evol. 8, 4619–4630, 10.1002/ece3.3979 (2018).

28. Goossens, K. V. et al. Molecular mechanism of flocculation self-recognition in yeast and its role in mating and survival. mBio 6, 1–16, 10.1128/mBio.00427-15 (2015).

29. Hamilton, W. D. The genetical evolution of social behaviour. I. J. Theor. Biol. 7, 1–16, 10.1016/0022-5193(64)90038-4 (1964).

30. Queller, D. C., Ponte, E., Bozzaro, S. & Strassmann, J. E. Single-gene greenbeard effects in the social amoeba Dictyostelium discoideum. Science 299, 105–106, 10.1126/science.1077742 (2003).

31. Foty, R. A. & Steinberg, M. S. The differential adhesion hypothesis: A direct evaluation. Dev. Biol. 278, 255–263, 10.1016/j.ydbio.2004.11.012 (2005).

32. Nowak, M. A. Five rules for the evolution of cooperation. Science 314, 1560–1563, 10.1126/science.1133755 (2006).

33. Nadell, C. D., Foster, K. R. & Xavier, J. B. Emergence of spatial structure in cell groups and the evolution of cooperation. PLoS Comput. Biol. 6, e1000716, 10.1371/journal.pcbi.1000716 (2010).

34. Drescher, K., Nadell, C. D., Stone, H. A., Wingreen, N. S. & Bassler, B. L. Solutions to the public goods dilemma in bacterial biofilms. Curr. Biol. 24, 50–55, 10.1016/j.cub.2013.10.030 (2014).

35. Boraas, M. E., Seale, D. B. & Boxhorn, J. E. Phagotrophy by flagellate selects for colonial prey: A possible origin of multicellularity. Evol. Ecol. 12, 153–164, 10.1023/A:1006527528063 (1998).

36. Staps, M., van Gestel, J. & Tarnita, C. E. Emergence of diverse life cycles and life histories at the origin of multicellularity. Nat. Ecol. Evol. 3, 1197–1205, 10.1038/s41559-019-0940-0 (2019).

37. De Vargas Roditi, L., Boyle, K. E. & Xavier, J. B. Multilevel selection analysis of a microbial social trait. Mol. Syst. Biol. 9, 684, 10.1038/msb.2013.42 (2013).

38. Damore, J. A. & Gore, J. Understanding microbial cooperation. J. Theor. Biol. 299, 31–41, 10.1016/j.jtbi.2011.03.008 (2012).

39. Denoth Lippuner, A., Julou, T. & Barral, Y. Budding yeast as a model organism to study the effects of age. FEMS Microbiol. Rev. 38, 300–325, 10.1111/1574-6976.12060 (2014).

40. Janssens, G. E. & Veenhoff, L. M. The natural variation in lifespans of single yeast cells is related to variation in cell size, ribosomal protein, and division time. PLoS ONE 11, e0167394, 10.1371/journal.pone.0167394 (2016).

41. Ross-Gillespie, A., Gardner, A., West, S. A. & Griffin, A. S. Frequency dependence and cooperation: Theory and a test with bacteria. Am. Nat. 170, 331–342, 10.1086/519860 (2007).

42. Healey, D., Axelrod, K. & Gore, J. Negative frequency-dependent interactions can underlie phenotypic heterogeneity in a clonal microbial population. Mol. Syst. Biol. 12, 877, 10.15252/msb.20167033 (2016).

43. Harrow, G. L. et al. Negative frequency-dependent selection and asymmetrical transformation stabilise multi-strain bacterial population structures. ISME J. 15, 1523–1538, 10.1038/s41396-020-00867-w (2021).

44. Avilés, L. Solving the freeloaders paradox: Genetic associations and frequency-dependent selection in the evolution of cooperation among nonrelatives. Proc. Natl. Acad. Sci. United States Am. 99, 14268–14273, 10.1073/pnas.212408299 (2002).

45. Fisher, R. M., Cornwallis, C. K. & West, S. A. Group formation, relatedness, and the evolution of multicellularity. Curr. Biol. 23, 1120–1125, 10.1016/j.cub.2013.05.004 (2013).

46. Pentz, J. T., Travisano, M. & Ratcliff, W. C. Clonal development is evolutionarily superior to aggregation in wild-collected Saccharomyces cerevisiae. In Artificial Life 14 - Proceedings of the 14th International Conference on the Synthesis and Simulation of Living Systems, ALIFE 2014, p550–554, 10.7551/978-0-262-32621-6-ch088 (MIT Press Journals, 2014).

47. Melbinger, A., Cremer, J. & Frey, E. The emergence of cooperation from a single mutant during microbial life cycles. J. Royal Soc. Interface 12, 10.1098/rsif.2015.0171 (2015). 1505.03558.

