## Supplementary Text for "Permissive aggregative group formation favors coexistence in yeast"

### Supplementary Information

### Methods

#### Yeast strains and growth

Yeast strains used are in Table 1. To fluorescently tag the flo1 deletion mutant, an yECitrine marker under the control of the TDH3 promoter was derived from KV1575[1] and transformed to the KV22 strain using the protocol of Gietz & Schiestl[2] (Table. 2). Transformants were selected on YPD 2% dextrose, 2% peptone, 1% yeast extract, 2% agar) supplemented with 100mg/l Hygromycin B and verified by PCR.

Yeast strains were inoculated from YPD ( plates stored at 4 °C in 5 mL YPD. Cultures were incubated at 30 °C at 200 RPM for three consecutive days. Afterwards, 100 µL of culture was inoculated in 5 mL YPG (2% galactose, 2% peptone, 1% yeast extract) for 2 consecutive days to induce potential flocculation. After growth and induction of *FLO1*, yeast cells were harvested by centrifugation at 1500 RPM for 5 minutes. The cells were resuspended in 3 mL of 200 mM EDTA to disrupt potential flocs. Subsequently cells were harvested by centrifugation and washed twice in milliQ.

#### Cantilever and AFM sample preparation

Single cell-force spectroscopy was based on the methods of El-Kirat-Chatel *et al.*[3] Tipless Pyrex-nitride probes (PNP-TR-TL, Nanoworld AG) were cleaned by UV-Ozone treatment for 15 min and were subsequently washed in distilled water. Cantilevers were subsequently functionalized by immersion in 10 mM Tris-HCl buffer (pH = 8.5) containing 4 $\mathrm{mg\,mL^{-1}}$ dopamince hydrochloride for 1 h. Concurrently, washed yeast cells were incubated with 20 µM FUN-1 cell (ThermoFisher) dye for 30 min in the dark, to validate cell viability. Cantilevers were thermally calibrated and were brought into contact with a single viable yeast cell for at least 3 min to allow attachment. FUN-1 stained yeast cells were inoculated on coverslips, identically polydopamine-functionalized as the cantilevers, and allowed to attach for 30 min. Afterwards, glass coverslips were extensively washed to remove unattached cells and kept in PBS at 4°C.

Single-cell force spectroscopy was performed on a Nanowizard III Bioscope-AFM (JPK instruments) in 10 mM Tris-HCL containing 200 µM $Ca^{2+}$. Probe velocity was varied between 0.5, 1 and 3 $\mathrm{\mu m\,s^{-1}}$ to estimate the linear bond friction using a Z-range of 3 µm. Per cell-cell pair a map of 16 by 16 force signatures with a scan size of 10 µm by 10 µm was obtained. These force signatures were further filtered on the contact point to retain only curves on top of the immobilized cell. From each curve the detachment force $F_d$ was obtained as the maximum adhesive force and the rupture distance $d_r$ as the maximal distance where there was a significant adhesive force.

#### Yeast flocculation assay

After washing, optical density was measured in 200 mM EDTA at 600 nm subsequently normalized to $OD_{600} = 1.7$ and $OD_{600} = 2.9$ for FLO+ and flo-, respectively. Normalized cultures FLO+ and flo- were mixed with various volume ratios to create various FLO+ frequencies. 0.5 mL of the mixed cultures was added to 4.5 mL milliQ or 10 mM $Ca^{2+}$ in falcon tubes. After addition of the cell cultures, the falcon tubes were gently inverted to homogenize their contents. After homogenization, 100 µL was sampled at approximately the center of liquid volume. Subsequently, the falcon tubes were agitated at 0, 100, 200 and 400 RPM for 5 min in an orbital shaker at 30°C and then placed upright for 5 min for floc sedimentation. At the end of sedimentation, 100 µL was sampled at the bottom of the falcon tube. 100 µL 200 mM EDTA was added to the sample before agitation and after agitation to disrupt potential flocs. Both samples were measured by flow cytometry and every condition was independently replicated three times.

#### Flow cytometry

Samples were measured with CytoFLEX Flow Cytometer (Beckman Coulter) using a 488 nm excitation laser. Yeast cells were distinguished from the background by gating the FSC and SSC ( Fig. S13A). Subsequently, doublets were removed from the cell population based on the area and height of the FSC signal (Fig. S13B). Finally, cells were classified on their cell type based on their fluorescent genomic constructs; respectively mCherry and yECitrine for FLO+ and flo- which were excited with a 488 nm laser and detected while using a 525/40 bandpass filter for yECitrine and 610/20 bandpass filter for mCherry (Fig. S13C). The cooperator frequency was calculated as the ratio of classified plus cells over the total number of cells. All gates were kept constant over all conditions and samples. Analysis was performed in CytExpert software (Beckman Coulter).

#### Individual cell-based model

Yeast cells are considered as spherical particles and were seeded randomly uniform in a cuboid with an elongated side axial to the shear flow $L_{\text{shear}}$. The cuboid has periodic boundary conditions in the shear direction and reflective boundary conditions in the perpendicular directions. The equation of motion for cell $i$ is the overdamped Langevin-type equation:

$$\lambda(v_i - v_i^f) = \sum_j F_{ij}^c + F^t(t),$$

where we used the friction constant $\lambda$, the contact force $F^c$ and a Brownian force $F^t$, with $\langle F^t(t)\rangle = 0$ and $\langle F^t(t)\cdot F^t(t')\rangle = 2\lambda k_B T\delta(t-t')$. $v^f$ denotes the velocity of the surrounding fluid, which is set by the linear shear field for cell $i$ with position $x_i$,

$$v_i^f = \dot{\gamma}(x_i \cdot \hat{e}_z)\,\hat{e}_y,$$

with $\dot{\gamma}$ the set shear rate. The friction coefficient $\lambda$ is approximated by Stokes' law for laminar flow over a sphere with radius $R$ in a fluid of viscosity $\eta$

$$\lambda = 6\pi\eta R\phi.$$

Here, $\phi$ is a simple correction for the shielding effect due to neighbours and is estimated as $\phi = \max(0, 1 - N_c/12)$, considering that a dense packing of spheres includes around 12 direct neighbours ($N_c$). The presence of this correction factor has no significant implications on the reported findings. The interaction force between cell $i$ and $j$ for a given overlap distance $d_{ij} = 2R - ||x_i - x_j||$ and contact normal unit vector $\hat{n}_{ij} = (x_i - x_j)/||x_i - x_j||$ is

$$F_{ij}^c = \begin{cases} \frac{4}{3}\hat{E}\sqrt{\hat{R}}\, d_{ij}^{3/2}\hat{n}_{ij} + \lambda_c(v_j - v_i) & \text{if } d_{ij} > 0, \\ -\frac{F_d}{d_r} d_{ij}\hat{n}_{ij} + \lambda_c(v_j - v_i) & \text{if } -d_r \le d_{ij} \le 0, \\ 0 & \text{if } d_{ij} < -d_r. \end{cases}$$

Cells experience a Hertz repulsive force for positive overlap distance, with effective Young's modulus $\hat{E} = \frac{E}{2(1-\nu)}$ and effective contact radius $\hat{R} = R/2$ (see Table 3). In the adhesive regime ($-d_r \le d_{ij} \le 0$), a linear adhesion force scales with the cell-cell overlap $d_{ij}$ until a maximum rupture distance $d_r$ is reached, as obtained by the AFM measurements, Fig. S1. In all regimens, the contact force includes a linear wet friction with friction constant $\lambda_c$. To solve the equations of motion, we make us of the combined overdamped equation

$$F = \Lambda v,$$

where $F$ is the total sum of forces on the particles and $\Lambda$ is the combined friction/resistance matrix that includes off-diagonal contributions for viscous cell-cell and cell-environment interactions. Note that the fluid drag force generates a contribution to both sides of this equation. The equations of motion are solved for the particle velocity using the conjugate gradient method. Next, particle positions are updated using an explicit Euler time integration scheme.

### Cluster size-based selection

The cooperative benefits of flocculation are attributed to the cluster size. Examples of cluster size-based advantages are protection against predators[4], cooperative metabolism[5], protection against chemical compounds[1], and increased sedimentation rates to escape unfavorable environments[6]. We opt for cluster size-based selection based on the relation between cluster size and the sedimentation velocity. For this we use an approximation of the Stokes' law using an effective diameter and effective density to correct for the fractal structure of the yeast clusters as proposed by Davis *et al.*[7]. The cluster sedimentation velocity $v_i$ in function of cluster size $C_i$ is

$$v_i = kC_i^{(1-f)/f}$$

where $k = a^{1/f}V_c(\rho_c - \rho)g/(3\pi\eta)$ with fractal prefactor $a$, yeast cell volume $V_c$, yeast cell density $\rho_c$, fluid density $\rho$, gravitational constant $g$ and fluid viscosity $\eta$ and fractal dimension $f$. we define survival probability based on the ratio of sedimentation velocities with the sedimentation velocity of a fully cooperative systems $(x_i = 1)$.

$$\begin{aligned} P(\text{survive}) &= 1 - \exp\left[-\frac{v_i}{\alpha\langle v_{(x_i=1)}\rangle}\right] \\ &= 1 - \exp\left[-\frac{C_i^{(f-1)/f}}{\alpha C_{(\infty|x_i=1)}^{(f-1)/f}}\right] \end{aligned}$$

where $\alpha$ is a variable parameter modulating the strength of selection. We opt for a fractal dimension of $f = 3$, however different fractal dimensions do not significantly alter our results (SI Fig. S9).

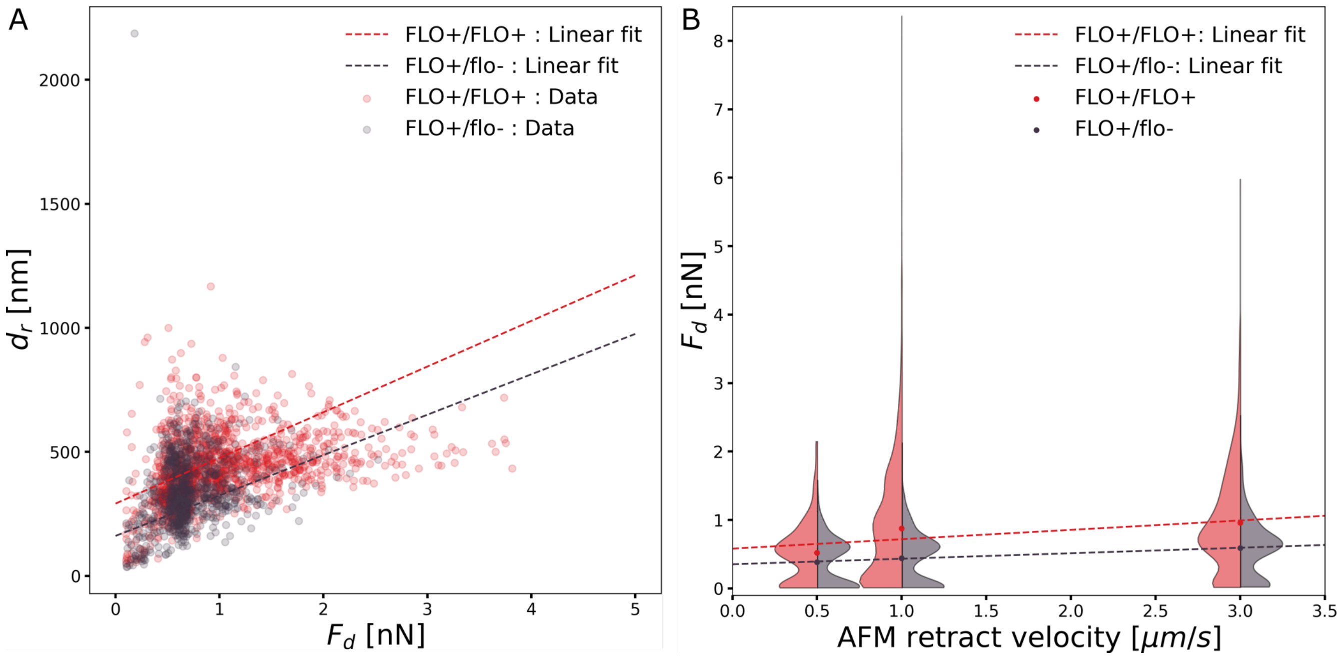


**Figure S1.** Further bond characterization of Flo1p flocculin (A) Experimental relationship between detachment force $F_d$ and bond rupture distance $d_r$ for homophilic FLO+/FLO+ interaction and heterophilic FLO+/flo- interactions. Measurements below single Flo1p-bond ($F_d < 100$ pN[3]) are not depicted. (B) Velocity dependency of the detachment force estimated by single cell-force spectroscopy with varying retraction velocity of the probe.

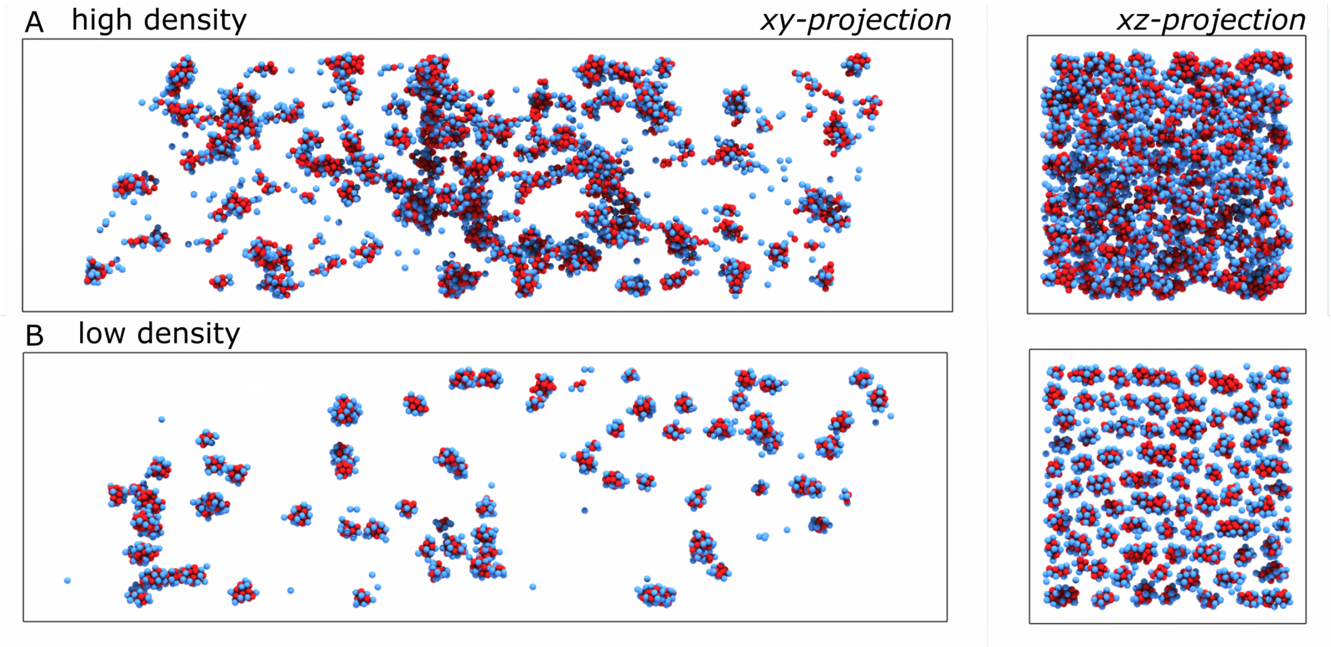


**Figure S2.** Comparison between high and low density. (A) In the high ('super-critical') density case, the xz-projected area integrated across a circular flow line is larger than one, resulting in a dynamic steady-state with continuous collisions. (B) At low ('sub-critical') density, this projected area is lower than one, and collisions become exceedingly rare after closed flow lines have been depleted of cells as indicated in the xz-projection.

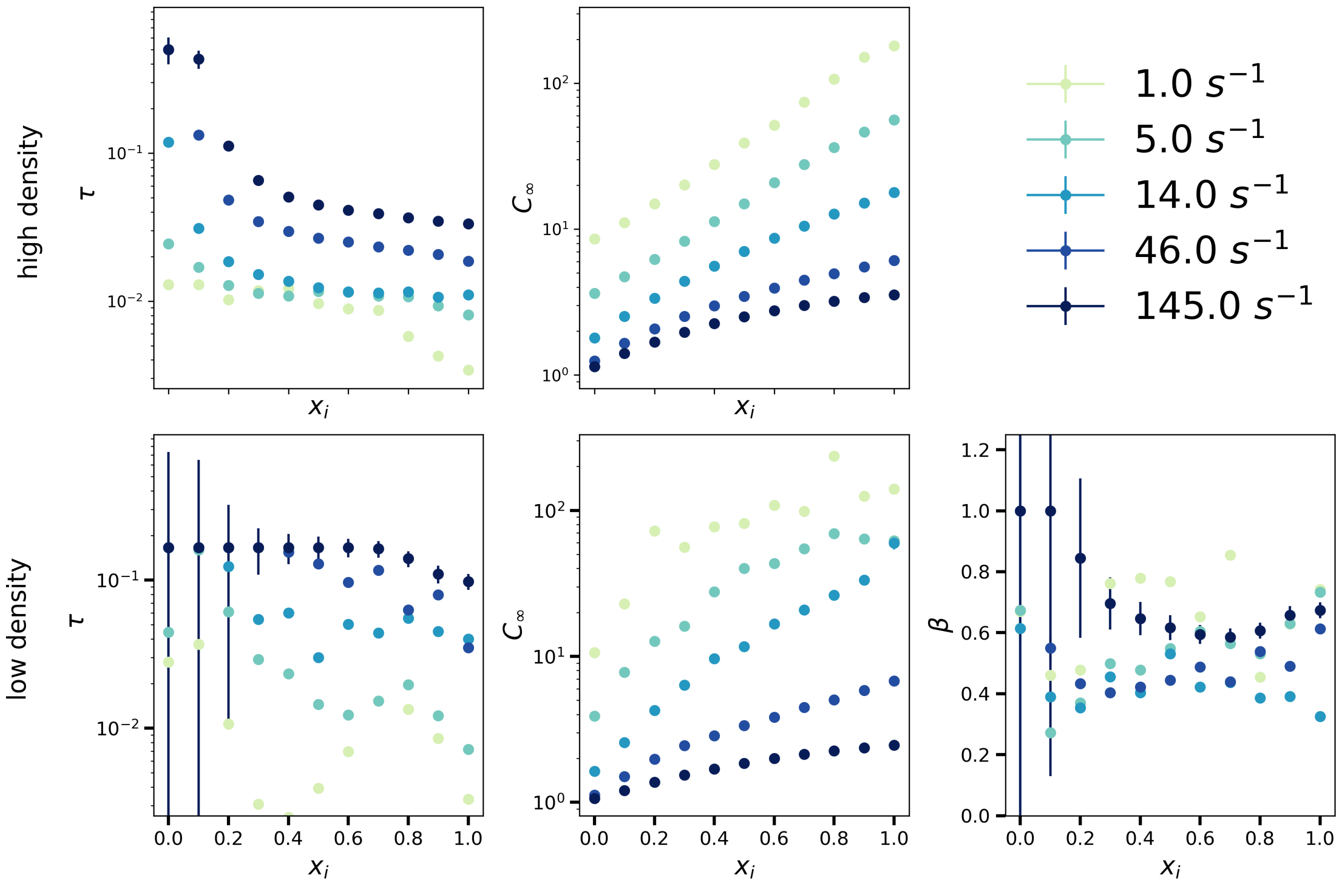


**Figure S3.** Parameter values of mean cluster size fits over time. $C(t) = C_\infty\,[1 - \exp(-t/\tau)]$ and a stretched exponential $C(t) = C_\infty[1 - \exp(-(t/\tau)^\beta)]$ fit for the high and the low density respectively. Note the high standard deviation for the stretched exponential at low $x_i$, indicative of over-fitting, since a regular exponential is sufficient for these data, rendering $b$ and $\beta$ interchangeable.

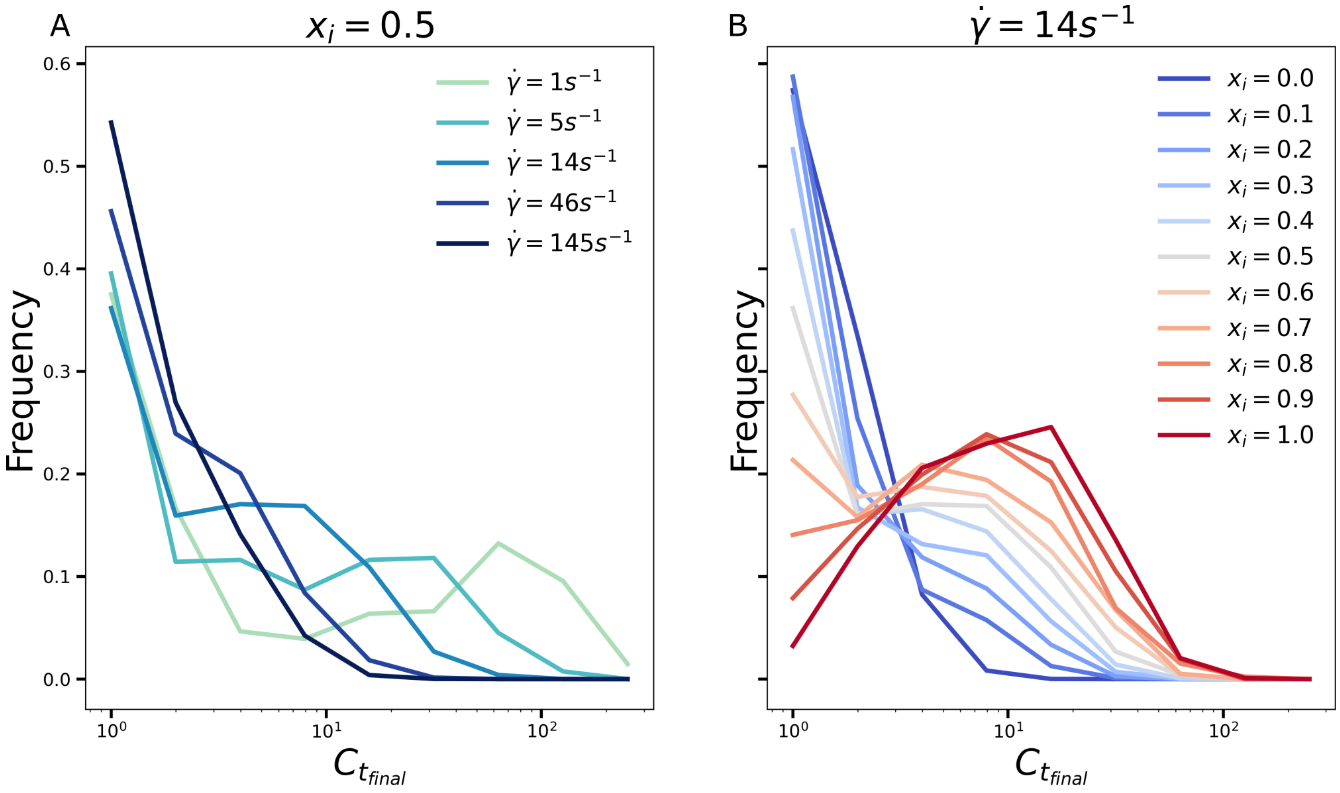


**Figure S4.** Cluster size distribution at the endpoint of flocculation $t_{final}$. (A) Cluster size distribution in function of shear rate $\dot{\gamma}$ with fixed cooperator frequency $x_i = 0.5$. (B) Cluster size distribution in function of cooperator frequency $x_i$ at shear rate $\dot{\gamma} = 14\,s^{-1}$.

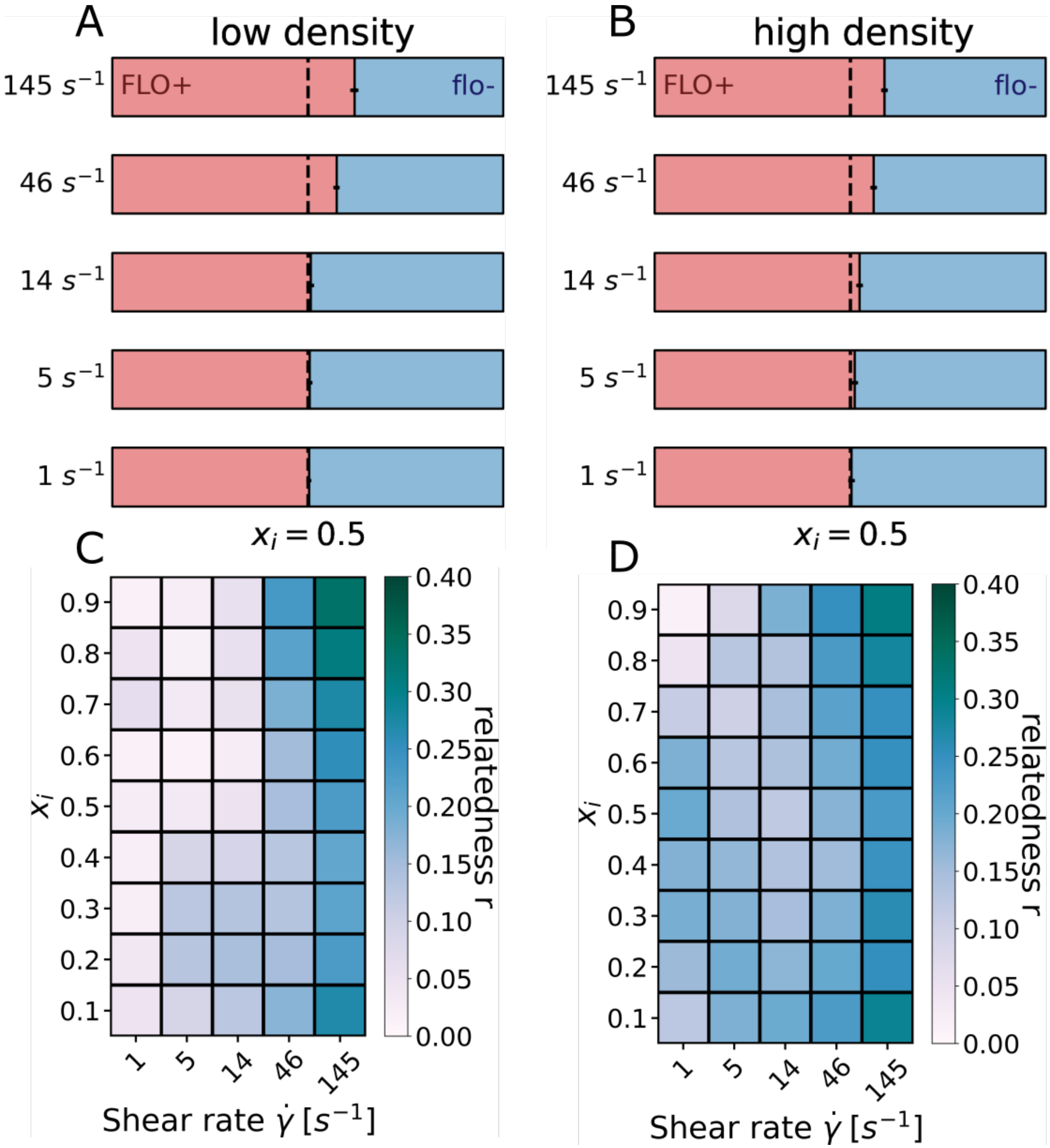


**Figure S5.** Kin recognition for high and low density. Cluster composition for clusters consisting of 2 or more cells for low (A) and high (B) density for increasing shear rates $\dot{\gamma}$. Cluster relatedness $r$ in function of shear rate $\dot{\gamma}$ and cooperator frequency $x_i$ for low (C) and high (D) density.

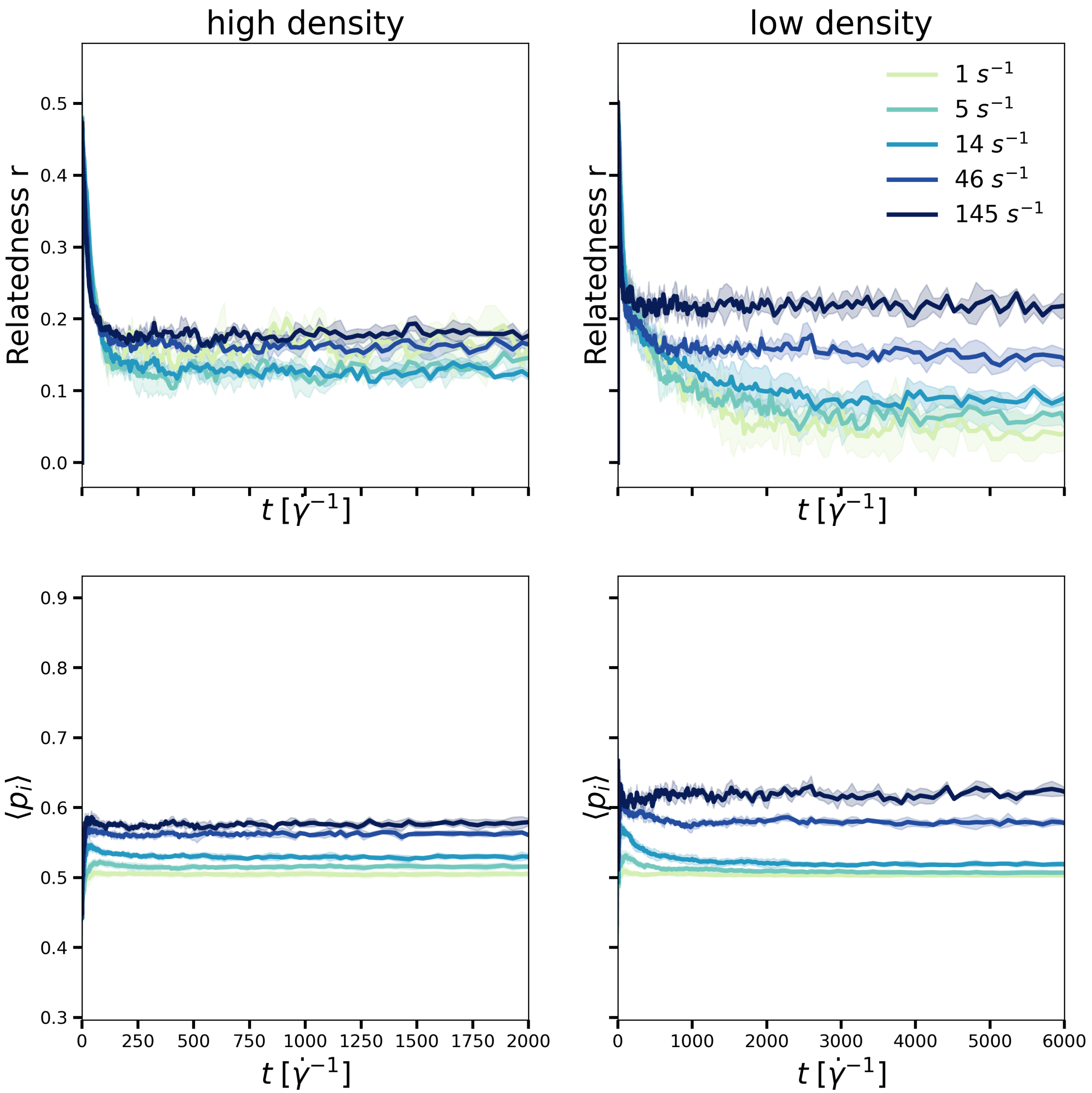


**Figure S6.** Exclusion dynamics for high and low density at cooperator frequency $x_i = 0.5$. (Top) Relatedness $r = (\langle p_i^2 \rangle - \langle p_i \rangle^2)/(\langle p_i \rangle - \langle p_i \rangle^2)$ with cluster composition $p_i$ over dimensionless time $\dot{\gamma}t$. (Bottom) Fraction of FLO+ cells in clusters larger than 2 cells. In case of P(FLO+) $> x_i$, there is enrichment of FLO+ cells in the cluster fraction.

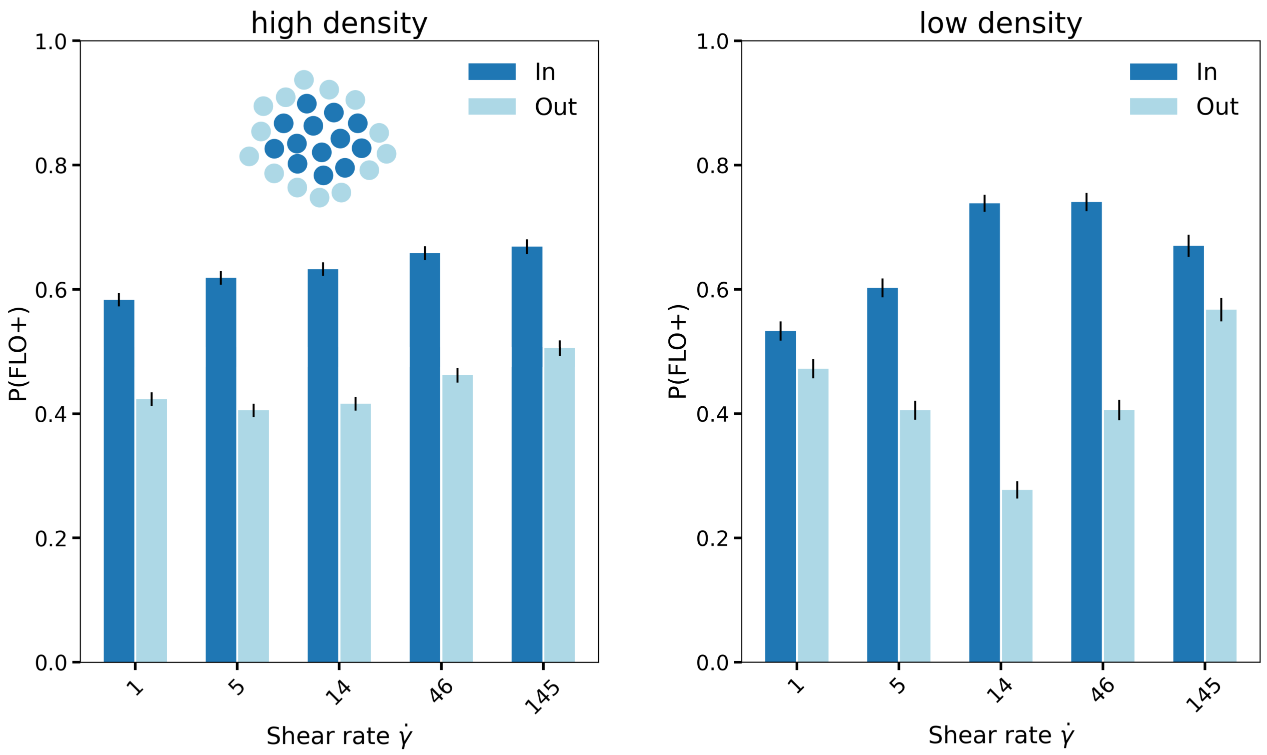


**Figure S7.** Radial assortment of cell types for cooperator frequency $x_i = 0.5$. The probability of cell being FLO+ $P(\text{FLO}+)$ was quantified separately for the 50% closest to the center of mass of the cluster (In) and the 50% cells farthest from the cluster center (Out) for high and low density.

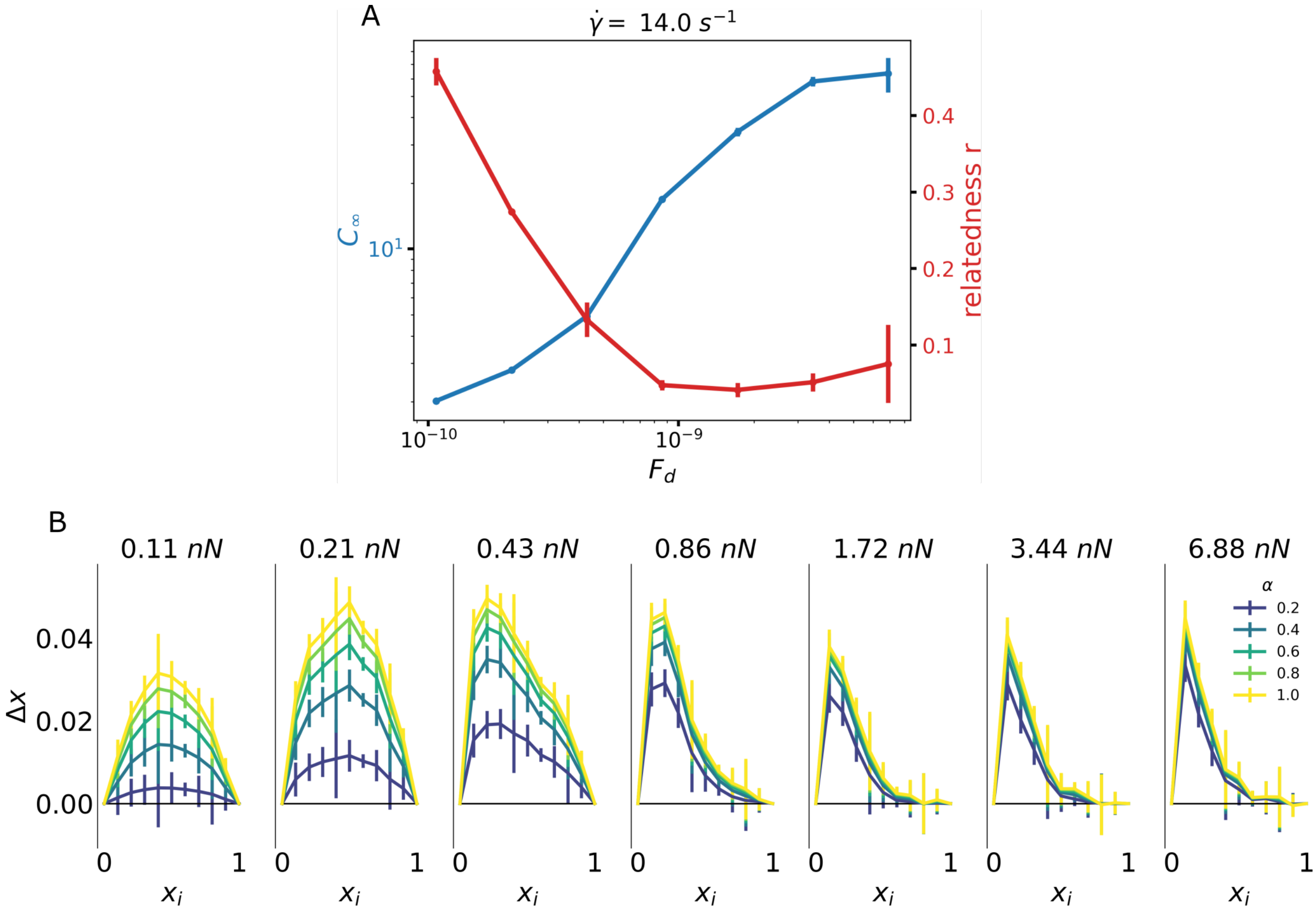


**Figure S8.** Effect of bond strength on kin recognition and cluster size. (A) Final mean cluster size $C_\infty$ and relatedness in function of FLO+/FLO+ detachment force $F_d$. $F_{+-}$ is scaled as $0.55F_{++}$, $F_{--}$ is kept constant at 0.06 nN. (B) Population drift curves with increasing adhesive strength for various selection strengths $\alpha$.

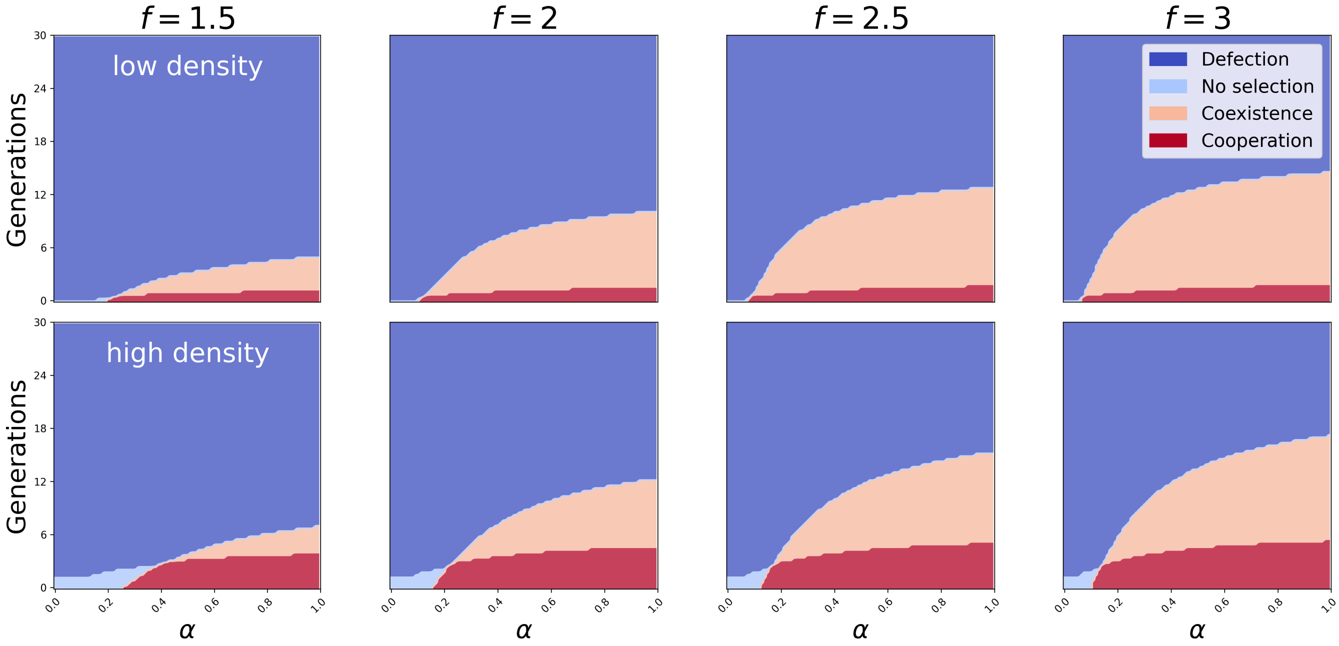


**Figure S9.** Effect of the fractal dimension $f$ on the predicted ESS for low and high density at $14\,s^{-1}$ for various selection strengths $\alpha$ and growth generations.

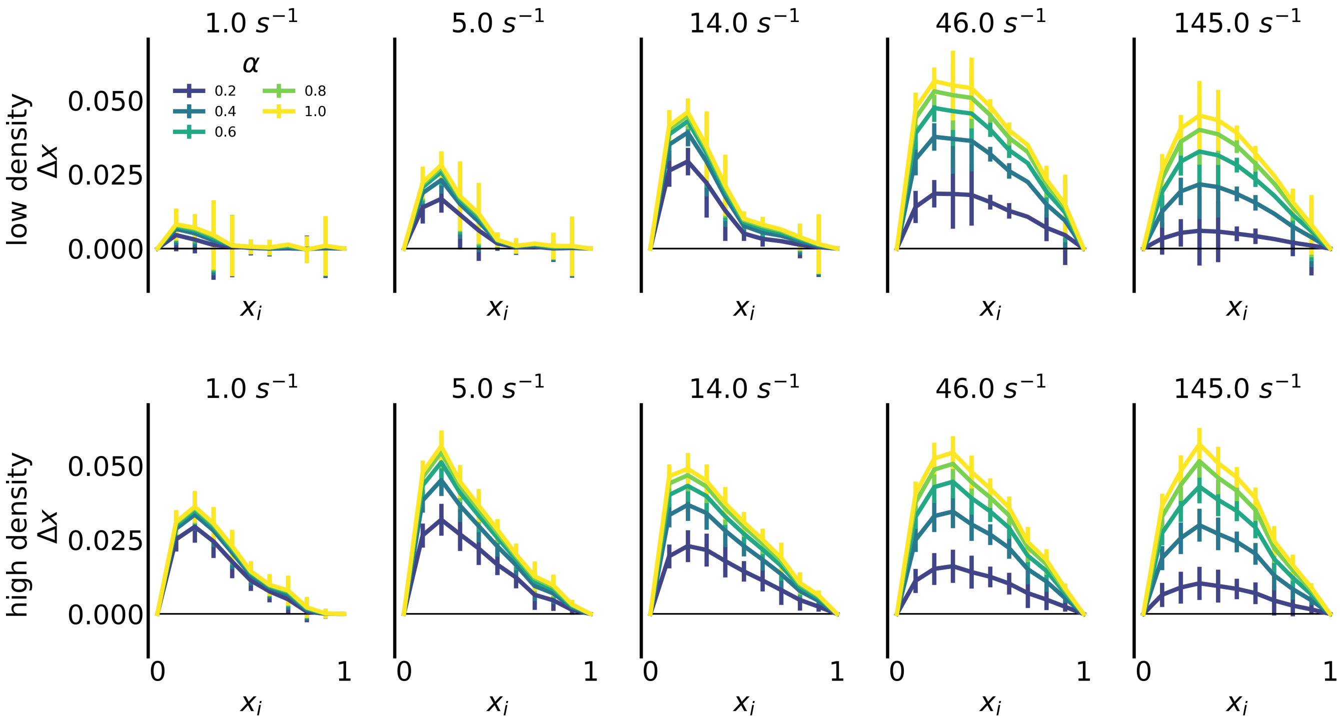


**Figure S10.** Evolutionary drift curves after flocculation, hence without any growth-associated fitness cost for Flo1p-production, for increasing selection strengths $\alpha$ for low and high density. The evolutionary drift $\Delta x$ given in function of the initial cooperator frequency $x_i$. Every condition shows a negative-frequency dependency in the evolutionary drift.

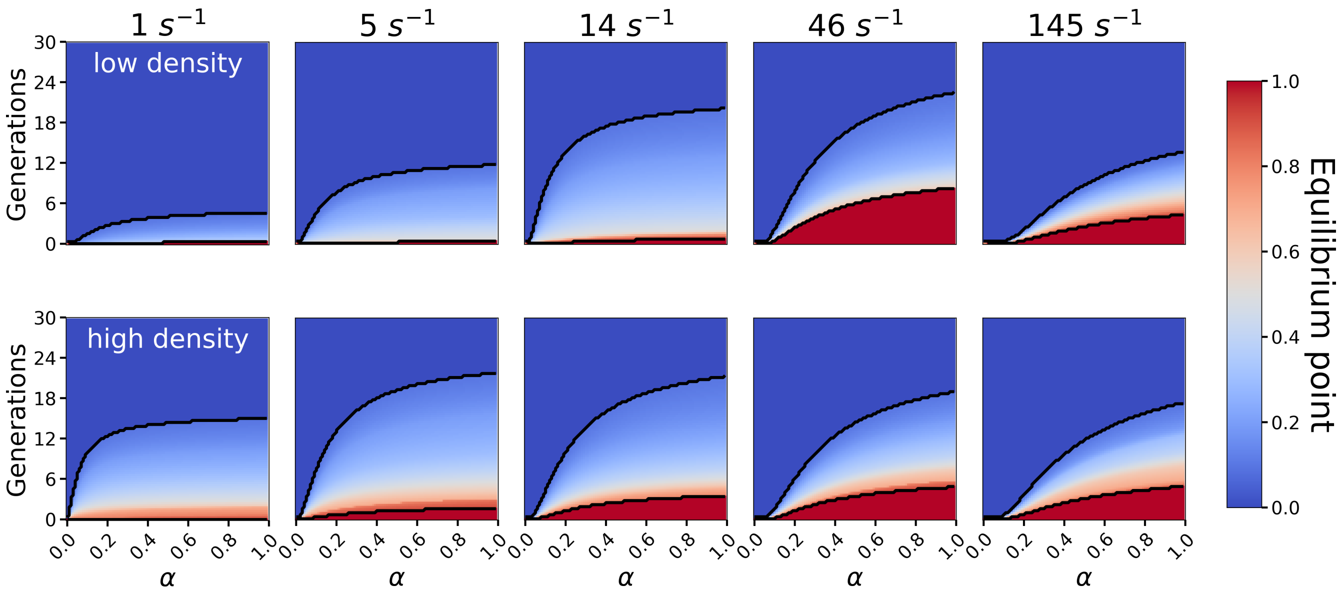


**Figure S11.** Evolutionary equilibrium point for high and low density. The equilibrium points are obtained by linearly interpolating between the data points and indicate the stable frequency of cooperators as predicted by the ESS. In case of cooperation, the equilibrium point equals one, in case of defection it equals zero. In case of coexistence the equilibrium is stable and lies between 0 and 1.

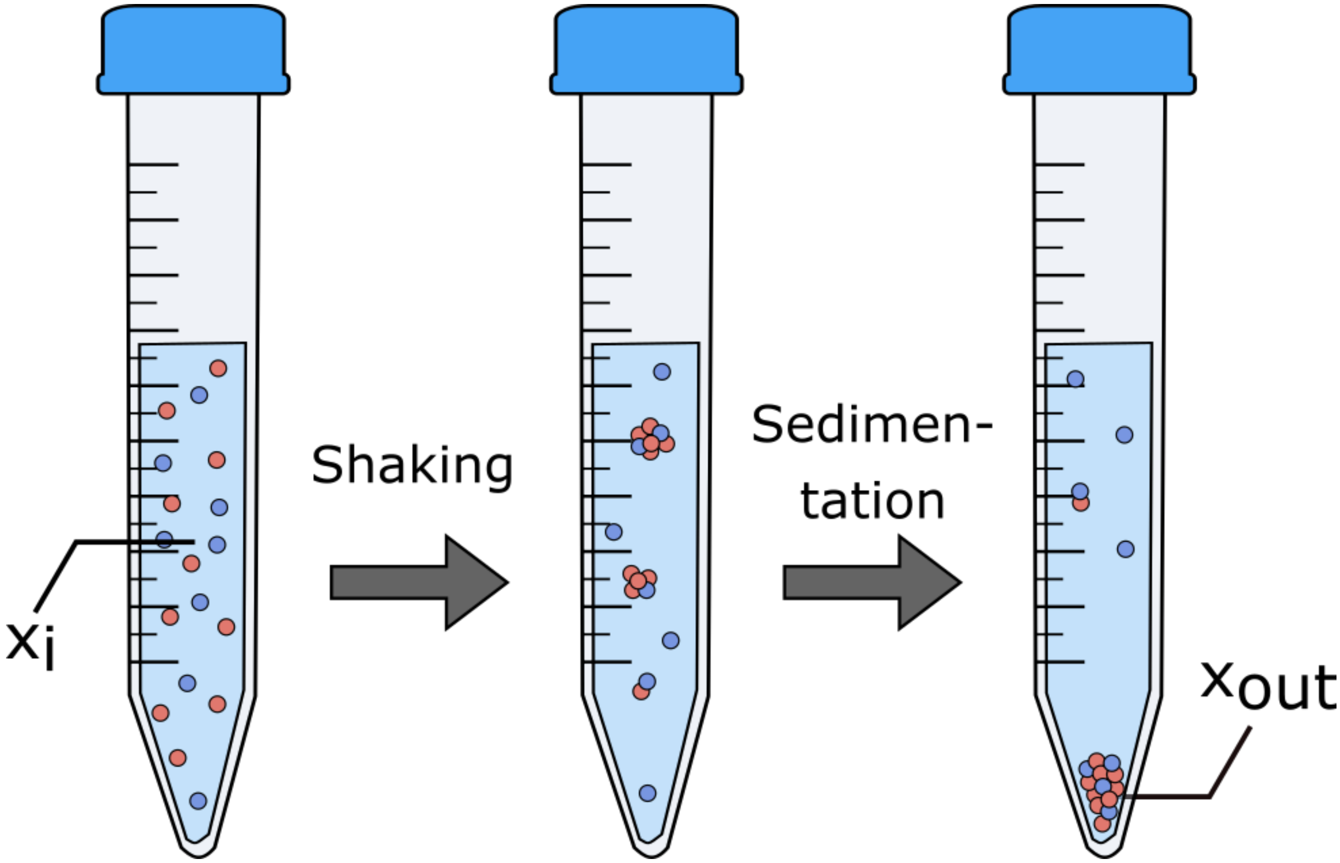


**Figure S12.** Schematic experimental set-up with drift determined by initial cooperator frequency $x_i$ and the cooperator frequency in the sedimented fraction $x_{\mathrm{out}}$.

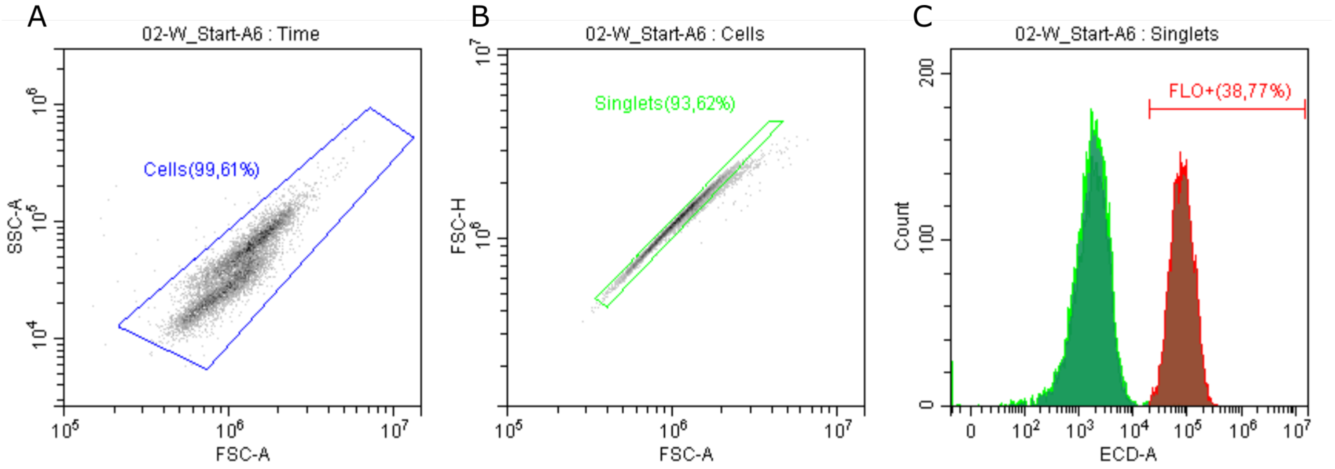


**Figure S13.** Processing of flow cytometry results (A) Cells are distinguished from the based on the area of the forward scattering signal (FSC-A) and side scattering signal. (B) Multiple cells simultaneously passing the detector are removed based on the height (FSC-H) and area (FSC-A) of the forward scatter. (C) Cell types are obtained by filtering on the excitation signal on the 525/40 bandpass filter for yECitrine and 610/20 bandpass filter for mCherry respectively, here shown for mCherry .

**Table 1.** List of yeast strains

| Strain | Relevant Genotype | Source |
|---|---|---|
| S288C (BY4742) | MAT$\alpha$ ; *his3*D1; *leu2*D0; *lys2*D0; *ura3*D0 | Brachmann *et al.* 1998[8] |
| KV210 | S288C BY4741 GAL1p-FLO1 | Verstrepen *et al.* 2005[9] |
| KV22 | S288C BY4741 *flo1::KANMX* | Smukalla *et al.* 2008[1] |
| KV1591 | KV210 TDH3p::mCherry-HYGB | Smukalla *et al.* 2008[1] |
| VK3876 | KV22 TDH3p::yECitrine-HYGB | This study |

**Table 2.** Oligonucleotides

| Oligo Name | Sequence |
|---|---|
| 147-HYG_Rv | TCGACAGACGTCGCGGTGAGTT |
| 1920-YRO-CONT-FW | CCTACCCGGTTGCCTCCAAAGCCCTTCTTT |
| 1923-YRO-CONT-RV | AACTAGGTTACATAACTGCCTATTGGCAAG |
| 3041-check-YRO2-f | GGCAACCATTCTCCAACATT |

**Table 3.** List of parameters used in the simulation

| Symbol | Description | Value | Unit | Source |
|---|---|---|---|---|
| $\dot{\gamma}$ | Shear rate | 1, 5, 14 ,46 ,145 | $s^{-1}$ | Varied |
| $dt$ | Timestep | $5\ 10^{-3}/\dot{\gamma}$ | s | Calibrated |
| $T$ | Temperature | 295 | K | Chosen |
| $R$ | Cell radius | 2.5 | µm | Milani *et al.* 1998[10] |
| $E$ | Cell Young's modulus | 30 | kPa | Calibrated |
| $\nu$ | Cell Poisson ratio | 0.4 | - | Stenson *et al.* 2011[11] |
| $\eta$ | viscosity of YPD | 6.9e-2 | Pa s | Calahorra *et al.* 2009[12] |
| $L_{\text{shear}}$ | Domain length shear direction | 600 | µm | Chosen |
| $L$ | Domain width and height | 200 | µm | Chosen |
| $\rho$ | Cell density | 1.66, 0.83 | cells $mL^{-1}$ | Varied |
| $\theta$ | Maximal conjugate gradient residual | 1e-4 | - | Calibrated |
| $F_{\text{d},++}$ | FLO+/FLO+ adhesion force | 0.86 | nN | Measured |
| $F_{\text{d},+-}$ | FLO+/flo- adhesion force | 0.47 | nN | Measured |
| $F_{\text{d},--}$ | flo-/flo- adhesion force | 0.06 | nN | Measured |
| $d_{\text{r},++}$ | FLO+/FLO+ rupture length | 450 | nm | Measured |
| $d_{\text{r},+-}$ | FLO+/flo- rupture length | 238 | nm | Measured |
| $d_{\text{r},--}$ | flo-/flo- rupture length | 50 | nm | Measured |
| $\lambda_{c,++}$ | FLO+/FLO+ bond friction | 84.0 | µN s $m^{-1}$ | Measured |
| $\lambda_{c,+-}$ | FLO+/flo- bond friction | 84.0 | µN s $m^{-1}$ | Measured |
| $\lambda_{c,--}$ | flo-/flo- bond friction | 3.0 | µN s $m^{-1}$ | Measured |

### References


1. Smukalla, S. *et al.* FLO1 Is a Variable Green Beard Gene that Drives Biofilm-like Cooperation in Budding Yeast. *Cell* **135**, 726–737, 10.1016/j.cell.2008.09.037 (2008).
2. Gietz, R. D. & Schiestl, R. H. High-efficiency yeast transformation using the LiAc/SS carrier DNA/PEG method. *Nat. Protoc.* **2**, 31–34, 10.1038/nprot.2007.13 (2007).
3. El-Kirat-Chatel, S. *et al.* Forces in yeast flocculation. *Nanoscale* **7**, 1760–1767, 10.1039/c4nr06315e (2015).
4. Kapsetaki, S. E. & West, S. A. The costs and benefits of multicellular group formation in algae*. *Evolution* **73**, 1296–1308, 10.1111/evo.13712 (2019).
5. Koschwanez, J. H., Foster, K. R. & Murray, A. W. Sucrose utilization in budding yeast as a model for the origin of undifferentiated multicellularity. *PLoS Biol.* **9**, e1001122, 10.1371/journal.pbio.1001122 (2011).
6. Goossens, K. V. *et al.* Molecular mechanism of flocculation self-recognition in yeast and its role in mating and survival. *mBio* **6**, 1–16, 10.1128/mBio.00427-15 (2015).
7. Davis, R. H. & Hunt, T. P. Modeling and Measurement of Yeast Flocculation. *Biotechnol. Prog.* **2**, 91–97, 10.1002/btpr.5420020208 (1986).
8. Brachmann, C. B. *et al.* Designer deletion strains derived from Saccharomyces cerevisiae S288C: A useful set of strains and plasmids for PCR-mediated gene disruption and other applications. *Yeast* **14**, 115–132, 10.1002/(SICI)1097-0061(19980130)14:2<115::AID-YEA204>3.0.CO;2-2 (1998).
9. Verstrepen, K. J., Jansen, A., Lewitter, F. & Fink, G. R. Intragenic tandem repeats generate functional variability. *Nat. Genet.* **37**, 986–990, 10.1038/ng1618 (2005).
10. Milani, M. *et al.* Differential two-color x-ray radiobiology of membrane/cytoplasm in yeast cells and lymphocytes. *Adv. Opt. Biophys.* **3256**, 195, 10.1117/12.307071 (1998).
11. Stenson, J. D., Hartley, P., Wang, C. & Thomas, C. R. Determining the mechanical properties of yeast cell walls. *Biotechnol. Prog.* **27**, 505–512, 10.1002/btpr.554 (2011).
12. Calahorra, M., Sánchez, N. S. & Peña, A. Activation of fermentation by salts in Debaryomyces hansenii. *FEMS Yeast Res.* **9**, 1293–1301, 10.1111/j.1567-1364.2009.00556.x (2009).
